# Asperous coordinates regenerative timing by regulating damage-induced WNT Signaling

**DOI:** 10.1101/2025.06.24.661394

**Authors:** Si Cave, Manashi Sonowal, Maksym Dankovskyy, Jordan Hieronymus, Chloe Van Hazel, Petra Fromme, Robin E. Harris

## Abstract

Tissue regeneration requires precise control of signaling pathways to direct proliferation, differentiation, and patterning. While early responses to injury are well characterized, how differentiation is coordinated during later stages remains unclear. Here, we identify Asperous (Aspr), an EGF-repeat protein, as a regeneration-specific regulator in *Drosophila* wing discs. Aspr is dispensable for wing development but is strongly induced within 24 hours post-injury. Maintaining *aspr* expression inhibits differentiation and alters reparative growth, while loss impairs regeneration. Structural and expression analyses show Aspr is an extracellular protein secreted in extracellular vesicles (EVs), where it co-localizes with the WNT ligand Wingless (Wg). We find Aspr regulates post-injury but not developmental Wg signaling, potentially by influencing its secretion or availability via EVs. These findings suggest Aspr regulates WNT activity to ensure proper timing of cell fate specification during regeneration, revealing a mechanism by which signaling dynamics are temporally controlled during tissue repair.

## Introduction

Tissue regeneration restores lost or damaged structures through coordinated reactivation of growth, differentiation, and patterning programs. Across different species and tissue types, this process frequently reuses developmental signaling pathways, including Wingless/Wnt (WNT), Bone Morphogenetic Protein (BMP), Notch, and JAK/STAT.^1–3^ These pathways drive key regenerative processes such as stem cell activation, proliferation, morphogenesis, and repatterning. Among them, WNT signaling is particularly well studied and broadly conserved,^4^ playing a pivotal role in regeneration of diverse tissues, including zebrafish fin and CNS,^5–9^ mouse and fly intestine,^10–13^ and planarian body axis.^14,15^ In many contexts, WNT ligands are rapidly induced after injury, activating canonical or non-canonical pathways to promote proliferative expansion.^4,16^ Moreover, WNT signaling is also required during the late stages of regeneration to re-establish spatial patterning cues, guiding the reconstruction of discrete tissue identities after proliferative growth.^4,16^ However, while the presence and function of WNTs and other pathways during regeneration are well documented, how their activity is precisely timed and regulated across different phases of regeneration remains poorly understood. This becomes critical for regenerating complex tissues through successive stages, which can require events like wound epithelium formation, development of a blastema, proliferative expansion, and tissue differentiation to occur sequentially.^1,17^ Each of these phases demand tight temporal control of signaling to maintain the correct progression of events, and understanding the mechanisms that coordinate such transitions remains an important challenge in the field.

The *Drosophila* wing imaginal disc provides an ideal model to dissect regeneration at high resolution, owing to its well-characterized developmental programs, advanced genetic tool kit and substantial ability to regenerate following injury.^18^ In addition to its historical use in studying growth and pattern formation,^19^ the wing disc has also emerged as a powerful system to identify genetic factors that regulate tissue regeneration.^18,20^ To better understand the events of injury induction and gene expression during regeneration, we previously developed a versatile genetic ablation system called DUAL Control.^21,22^ This system enables independent induction of cell death alongside spatially and temporally restricted manipulation of gene expression in surrounding tissues, allowing us to interrogate regeneration in a controlled and reproducible manner. Using this approach, we previously identified and characterized several different genetic and epigenetic factors that are required for proper restoration of the wing disc after injury.^22–25^

One such factor is *asperous* (*aspr*), a gene that encodes a putative extracellular EGF repeat–containing protein that is highly induced in early regenerating tissue. We initially identified *aspr* based on its proximity to a damage-responsive enhancer and its strong transcriptional upregulation following ablation,^22^ although its function in regeneration remained unclear. Here we have performed a comprehensive characterization of *aspr*, finding that it is not expressed during normal wing disc development and is dispensable for both viability and wing formation, yet its loss impairs regeneration, suggesting a regeneration-specific role. We show that *aspr* is present only transiently during the early phase of regeneration, and that prolonging its expression disrupts proper regenerative growth and delays the onset of late-stage patterning. Mechanistically, we find that *aspr* interferes with the reactivation of several patterning genes normally induced late in regeneration downstream of *Drosophila Wnt1*, *wingless* (*wg*), indicating *aspr* normally modulates WNT signaling. Further analysis reveals that the Aspr protein is secreted, and associates with extracellular vesicle-like structures, which colocalize with Wg protein explicitly during regeneration, suggesting it could potentially act as an extracellular regulator of WNT ligand distribution or availability in this context. Although vesicle-mediated transport has been proposed as a mechanism for WNT ligand trafficking during development,^26–29^ its role in regeneration has not been previously demonstrated. Thus, our work identifies a novel, regeneration-specific modulator of WNT signaling and highlights the critical role of regulating extracellular signaling to precisely coordinate gene expression timing necessary for successful tissue regeneration. Given the conserved role of WNT signaling in regeneration across diverse species,^16^ these findings have broad implications for understanding how signaling transitions are coordinated during regeneration.

## Results

### Damage-induced and developmental *aspr* are regulated by distinct mechanisms

Our previous work to characterize epigenetic and transcriptomic changes associated with regeneration identified *aspr* (previously *CG9572*),^22,25^ which RNA-seq showed to be one of the most strongly induced genes in regenerating discs.^25^ Other groups have shown that *aspr* is damage-responsive and associated with blastema cells.^30,31^ Notably, these studies also show *aspr* is absent from undamaged pouch tissue suggesting a regeneration-specific role, although this role remains unclear. To better understand its function, we visualized *aspr* expression after damage using HCR fluorescence *in situ* for transcript detection and an anti-Aspr antibody generated for this study. In undamaged early L3 discs (84 h after egg deposition, AED), *aspr* is expressed in the proximal notum and weakly above the dorsal hinge but is absent from the pouch (Figure 1A). This pattern persists in late L3 (108 h AED, Figure 1G). Unfortunately, the anti-Aspr antibody we developed exhibits low sensitivity and only weakly detects Aspr protein (Figure S1A and S1A’). Therefore, we relied on transcript detection for subsequent analyses. Using the DUAL Control (DC) ablation system^21^ to activate JNK-mediated apoptosis via activated *hempiterous* in the distal pouch (DC^hepCA^), we found *aspr* is upregulated in blastema cells by 12 h after heat shock (AHS), maintained through 18-24 h, and largely absent by 36 h (Figures 1B–E, quantified in 1F). By contrast, late L3 discs show only weak *aspr* upregulation following ablation (Figure 1H). We also tested necrotic cell death using DC^gluR1^.^23,24^ Similar to apoptosis, necrosis triggers *aspr* induction around the wound in early L3, peaking at 18 h and diminishing by 36 h (Figures 1I–J). Notably, however, necrosis causes *aspr* to occur with a more punctate, potentially nuclear, appearance (Figure 1I), likely reflecting the distinct responses to necrotic injury we previously characterized.^23,24^ Late L3 discs show minimal *aspr* expression post-necrosis (Figure 1K).

**Figure 1.**
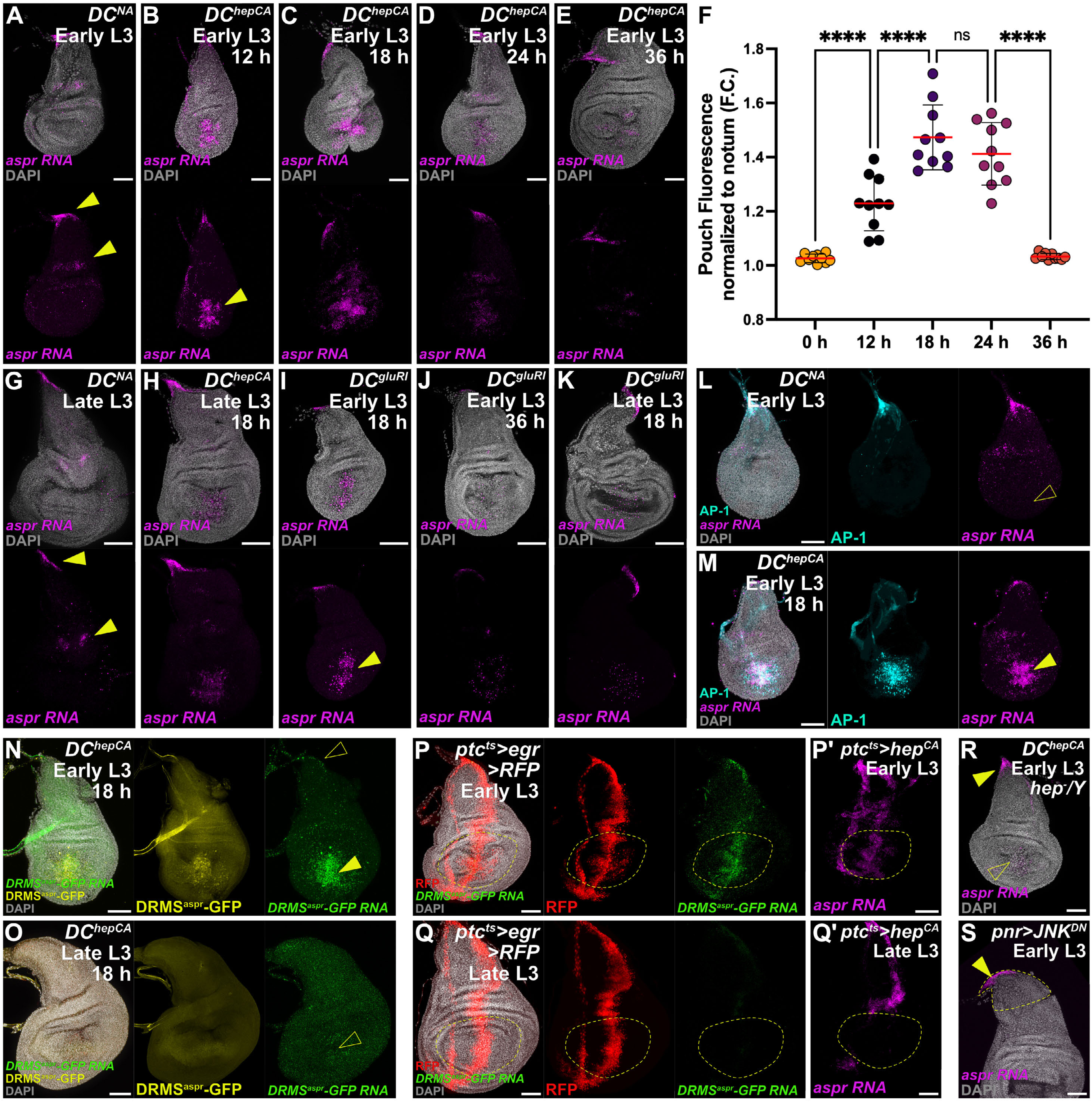
*aspr* is a damage-responsive gene regulated by a DRMS enhancer. (**A–E**) Time course of *aspr* RNA (magenta) in early L3 wing discs. Unablated control (*DC^NA^*) is shown in (A). Following apoptotic ablation (*DC^hepCA^*), *aspr* is strongly induced by 12 h AHS (arrowhead in B), maintained through 24 h, and reduced by 36 h (C–E). Nuclei are stained with DAPI (gray). Notum and dorsal hinge developmental expression indicated (yellow arrowheads in A), and damage-responsive expression in ablated disc (yellow arrowhead in B) (**F**) Quantification of *aspr* fluorescence measured in the pouch normalized to the notum. Statistics: one-way ANOVA; sample size = 10 discs per time point; ns = not significant; **** p < 0.0001. (**G–H**) *aspr* RNA in late L3 unablated disc (G) and ablated disc (H) showing reduced damage-induced expression in late L3. (**I–K**) Necrotic ablation (*DC^gluR1^*) in early L3 induces robust *aspr* at 18 h (I, arrowhead) that diminishes by 36 h (J); expression is weaker following late L3 ablation (K). (**L–M**) AP-1-RFP (cyan) overlaps with damage-induced *aspr* in DC^hepCA^ ablated discs (M, arrowhead) but not in unablated controls (L, open arrowhead). (**N–O**) RNA *in situ* for *DRMS^aspr^-GFP* (yellow) and *GFP* RNA (green) shows overlap in early L3 ablated discs (N, arrowhead) but weak expression in late L3 (O, open arrowhead); *GFP* RNA is absent from the notum (open arrowhead in N). (**P–Q**′) *ptc>egr* (P,Q) and *ptc>hepCA* (P′,Q′) ablation induce strong early L3 *DRMS^aspr^-GFP* and *aspr* RNA expression (P,P′) that diminishes in late L3 (Q,Q′). Dotted outlines indicate the pouch. (**R**) DC^hepCA^ ablation in *hep^-^/Y* null hemizygous mutants shows reduced *aspr* in the ablated pouch (open arrowhead) but preserved developmental expression (arrowhead). (**S**) Blocking JNK in the notum with *pnr>JNK^DN^* does not suppress developmental *aspr* expression (arrowhead). All scale bars = 50 μm unless indicated. See also Figure S1. Full genotypes listed in Supplementary Genotypes.

Beyond genetic injury, *aspr* is also expressed in neoplastic tumors (*sd>lglRNAi*, Figure S1B–D) and soon after physical wounding (Figure S1E), supporting its role as a generalized early stress-response gene. Similarly, in other imaginal discs (leg, haltere, eye), *aspr* is mostly undetectable in undamaged tissue but becomes induced upon ablation (Figures S1F–H). As with many genes involved in regeneration, *aspr* appears to be regulated by a Damage-Responsive, Maturity-Silenced (DRMS) enhancer, a class of regulatory elements that are activated by JNK and JAK/STAT signaling but epigenetically silenced in mature discs.^22,25,32^ We previously identified a DRMS enhancer upstream of *aspr* that becomes accessible after damage in early L3 but not late L3 discs.^22^ A reporter for this region (*DRMS^aspr^-GFP*) contains AP-1 and Stat92E binding sites (Figure S1I), overlaps with JNK activity (*AP-1-RFP*, Figures 1L–M), and mirrors *aspr* expression in early L3, while being only weakly active in late L3 (Figures 1N–O). This behavior is consistent with the DRMS regulating *aspr* via JNK in the context of damage. Since regenerative ability varies across the disc,^33^ we explored the regionality of DRMS^aspr^ activity by ablating tissue along the anterior/posterior (A/P) boundary using *ptc>hep^CA^*or *ptc>egr*. This domain includes the notum, hinge, and pouch (Figure 1P and Q), all of which activate JNK after damage.^33^ While only the pouch typically activates regeneration-associated genes, *aspr* is induced along the full stripe in early L3 discs (Figures 1P–P’), including non-regenerating notum tissue, although *DRMS^aspr^-GFP* is primarily activated in the pouch (Figure 1P). In late L3, *aspr* is only induced outside the pouch (Figure 1Q–Q’), while the *DRMS^aspr^-GFP* reporter is entirely inactive (Figure 1Q). Thus, damage-induced *aspr* is likely regulated by the DRMS^aspr^ in the regeneration-capable pouch, while damage-responsive enhancers that are not silenced with maturity likely drive *aspr* expression outside of this region.

Interestingly, we noted that developmental *aspr* expression in the notum overlaps with JNK activity that is required for thorax closure during pupariation (Figure 1A).^34^ In *hep^-^* mutants, damage-induced *aspr* is diminished, while developmental expression persists (Figure 1R). Similarly, expression of a dominant-negative JNK in the notum (*pnr>hep^DN^*) fails to suppress *aspr* in the notum (Figures 1S). Since the DRMS^aspr^ reporter is inactive in the notum of undamaged discs (Figure 1N), together these observations suggest that developmental and regenerative *aspr* are regulated by distinct mechanisms.

### Loss of *aspr* blocks regeneration but does not affect normal development of the wing disc

As *aspr* is strongly upregulated after injury but shows limited developmental expression, we tested whether its removal affects either context. The DC system allows heat shock-induced, flip-out-mediated expression of UAS-transgenes specifically in the pouch, driven by *DVE>>GAL4* during and after damage.^21^ We identified an RNAi construct that strongly rescues damage induced *aspr* in the pouch (Figures 2A–B) A second RNAi line (*TRiP.HMJ22471*) was tested, but we found it to be less effective and did not use it further. The knockdown of *aspr* in undamaged discs has no visible effect on wing development (Figure 2C), consistent with its minimal expression in normally developing discs (Figure 1A). However, knockdown during regeneration significantly impairs recovery in both early and late L3, shown by wing scoring (Figure 2D–G) and adult wing area (Figure 2H). The loss of *aspr* does not alter damage-induced JNK signaling, shown by JNK targets *wg* and *Mmp1*, or cell death caused by DC^hepCA^ ablation (Figures 2I–J and M–N). Although these data do not indicate whether Aspr is a direct target of JNK signaling, or a consequence of cell death, it does support the hypothesis that Aspr is downstream of, and regulated by, a JNK-responsive DRMS enhancer (Figure 1). Beyond these observations, the knockdown of *aspr* does not immediately indicate its function, though it is clearly essential for regeneration and mostly dispensable for normal development.

**Figure 2.**
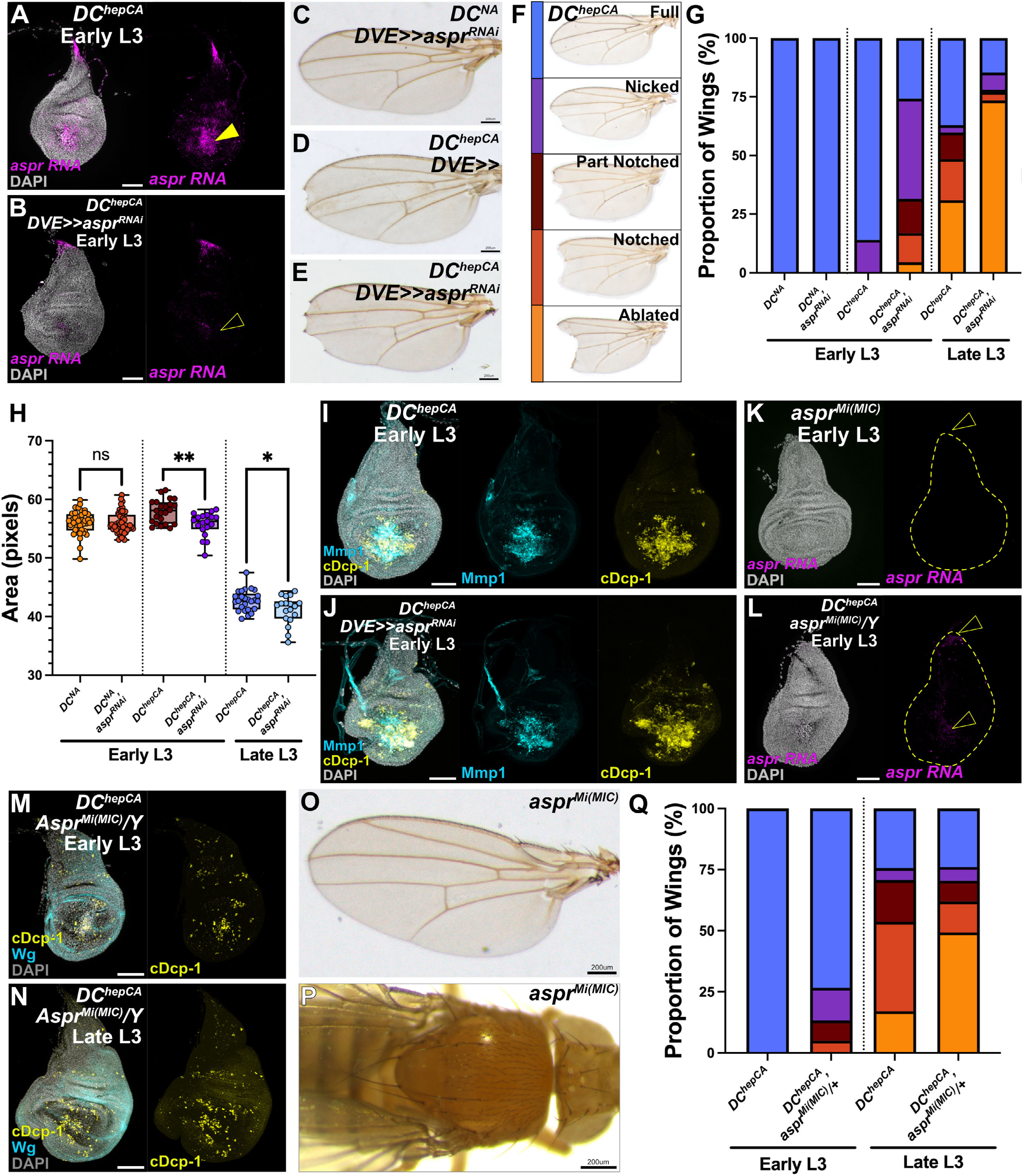
*aspr* is required for regeneration but appears dispensable for normal development. (**A–B**) *aspr* RNA (magenta) in early L3 discs imaged 18 h AHS after DC^hepCA^ ablation. Control (A) shows strong damage-induced expression (arrowhead); knockdown of *aspr* with *DVE>>aspr^RNAi^* (B) strongly reduces damage-induced *aspr* (open arrowhead). Nuclei = DAPI (gray). (**C–E**) Representative adult wing outcomes: (C) knockdown of *aspr* with DVE>>aspr^RNAi^ in undamaged discs (DC^NA^) has no effect; (D) control ablated (*DC^hepCA^, DVE>>)* discs have wild-type recovery; (E) knockdown of *aspr* with DVE>>aspr^RNAi^ during regeneration results in impaired recovery. (**F**) Definition of regeneration scoring categories used for adult wings following ablation with DC^hepCA^. (**G**) Regeneration scoring plotted as proportion of wings in each category for undamaged (*DC^NA^*) or ablated (*DC^hepCA^*) discs in early or late L3, with or without *aspr* knockdown. *aspr* knockdown reduces regeneration at both stages. Sample sizes: DC^NA^: *y^RNAi^*n = 134, *aspr^RNAi^* n = 132; Early L3: *y^RNAi^* n = 50, *aspr^RNAi^* n = 89; Late L3: *y^RNAi^* n = 97, *aspr^RNAi^* n = 109. (**H**) Quantification of adult wing area (box plots, median and quartiles) from wings in (G). Knockdown of *aspr* during development (*DC^NA^*) does not affect adult wing size; knockdown following ablation significantly reduces wing area. Statistics: one-way ANOVA. Sample sizes: *DC^NA^: y^RNAi^* n = 39, *aspr^RNAi^* n = 52; Early L3: *y^RNAi^* n = 23, *aspr^RNAi^* n = 19; Late L3: *y^RNAi^* n = 25, *aspr^RNAi^* n = 18. ns = not significant; * p = 0.0266; ** p = 0.0013. (**I–J**) Ablated (*DC^hepCA^*) early L3 discs stained for Mmp1 (cyan) and cleaved Dcp-1 (cDcp-1, yellow). *aspr* knockdown does not alter Mmp1 or cDcp-1 levels relative to control. (**K–L**) Early L3 discs from *aspr^Mi(MIC)^* mutant larvae: homozygous mutant lacks developmental *aspr* expression in the notum (K, open arrowhead) and ablated *aspr^Mi(MIC)^/Y* discs lack both developmental and damage-induced *aspr* (L, open arrowheads). (**M–N**) Ablated *aspr^Mi(MIC^*^)^/*Y* discs stained for Wg (cyan) and cDcp-1 (yellow) show that damage-induced Wg and apoptosis are unaffected in mutant discs at early (M) and late (N) L3. (**O–P**) Adult wing (O) and thorax (P) from *aspr^Mi(MIC)^* adults show no obvious developmental defects. (**Q**) Regeneration scoring for *w^1118^* control and *aspr^Mi(MIC)^/+* heterozygotes following ablation (*DC^hepCA^*). Heterozygotes show reduced regeneration at both stages. Sample sizes: Early L3: *w^1118^* n = 18, *aspr^Mi(MIC)^/+* n = 60; Late L3: *w^1118^* n = 41, *aspr^Mi(MIC)^/+* n = 66. All scale bars = 50 μm unless otherwise specified. See also Figure S2. Full genotypes in Supplementary Genotypes.

To confirm this, we tested the *aspr^MI02471^* allele, a *Mi[MIC]* insertion that likely represents a transcriptional null.^22^ HCR *in situ* confirms that hemizygous males and homozygous females lacked detectable *aspr* transcripts in the wing disc, both in undamaged and damaged conditions (Figures 2K–L). These mutants displayed no overt developmental phenotypes, including in wing patterning or size (Figure 2O). Moreover, while *aspr* is normally expressed in the notum, where JNK and other signals govern thoracic patterning and bristle formation, these structures were unaffected in mutants (Figure 2P). We did observe a modest but significant developmental delay in *aspr^MI02471^* larvae, which is more pronounced in homozygous females (Figures S2A and C). This delay is not seen with RNAi knockdown in the wing (Figure S2B), suggesting there is either a role for *aspr* in developmental timing outside of the wing, or an unrelated mutation in the *Mi[MIC]* line. Importantly, as in the knockdown experiments, *aspr^MI02471^* mutants showed reduced regenerative capacity in both early and late L3 discs (Figure 2Q). Thus, although the specific function of *aspr* remains unclear, these data confirm that it is required for regeneration but plays a minimal role in normal development.

### Maintaining *aspr* expression impacts repatterning during regeneration

*aspr* is normally upregulated only during the first 24 h of repair (Figures 1A–1F), and while loss-of-function analyses show it is required for regeneration, they do not clarify its role. Since *aspr* seems mostly dispensable for normal development but essential for early regeneration (Figure 2), we investigated its function via ectopic expression. We previously used a UAS-driven multi-tagged construct (*UAS-aspr-FLAG.3xHA*) to perform preliminary investigations of its effects,^22^ but this causes ectopic vein tissue in undamaged discs, a phenotype that is not seen with the untagged version we generated (*UAS-aspr*) (Figures S3A–B). Additionally, the multi-tagged construct showed inconsistent transcript levels in discs (Figure S3C). To ensure reliable findings, we generated a new single HA-tagged transgene (*UAS-aspr-HA*) and expressed it using both non-ablating (DC^NA^) and ablating (DC^hepCA^) versions of DUAL Control. Expression of the untagged and single tagged was confirmed by HCR and/or anti-HA staining (Figures 3A–B’, S3D–E). Strikingly, we found that Aspr protein occurs with two distinct appearances: “cellular Aspr,” associated with apical membranes, and “punctate Aspr,” found as discrete foci at or above the apical surface (Figure 3A-A’ and Figure 7G–K). This appearance was seen when expressed in the notum, hinge and pouch regions of the disc (Figure 3B–B’), but was not observed with the FLAG.3xHA multiple-tagged construct (Figure S3F–F’). Despite these distinct appearances, ectopic Aspr expression during development in the pouch (*DVE>>GAL4*), or throughout the entire anterior or posterior disc (*ci-GAL4* or *hh-GAL4*) had no visible effects on disc proliferation, signaling pathways, or gene expression (Figures 3C–H and S6A–D), nor did it affect adult wing patterning or size (Figures 3I-K and S3G-H). This held true for both the single HA-tagged and untagged constructs (Figure S3B), consistent with the notion that Aspr has limited developmental impact, even when ectopically expressed.

**Figure 3.**
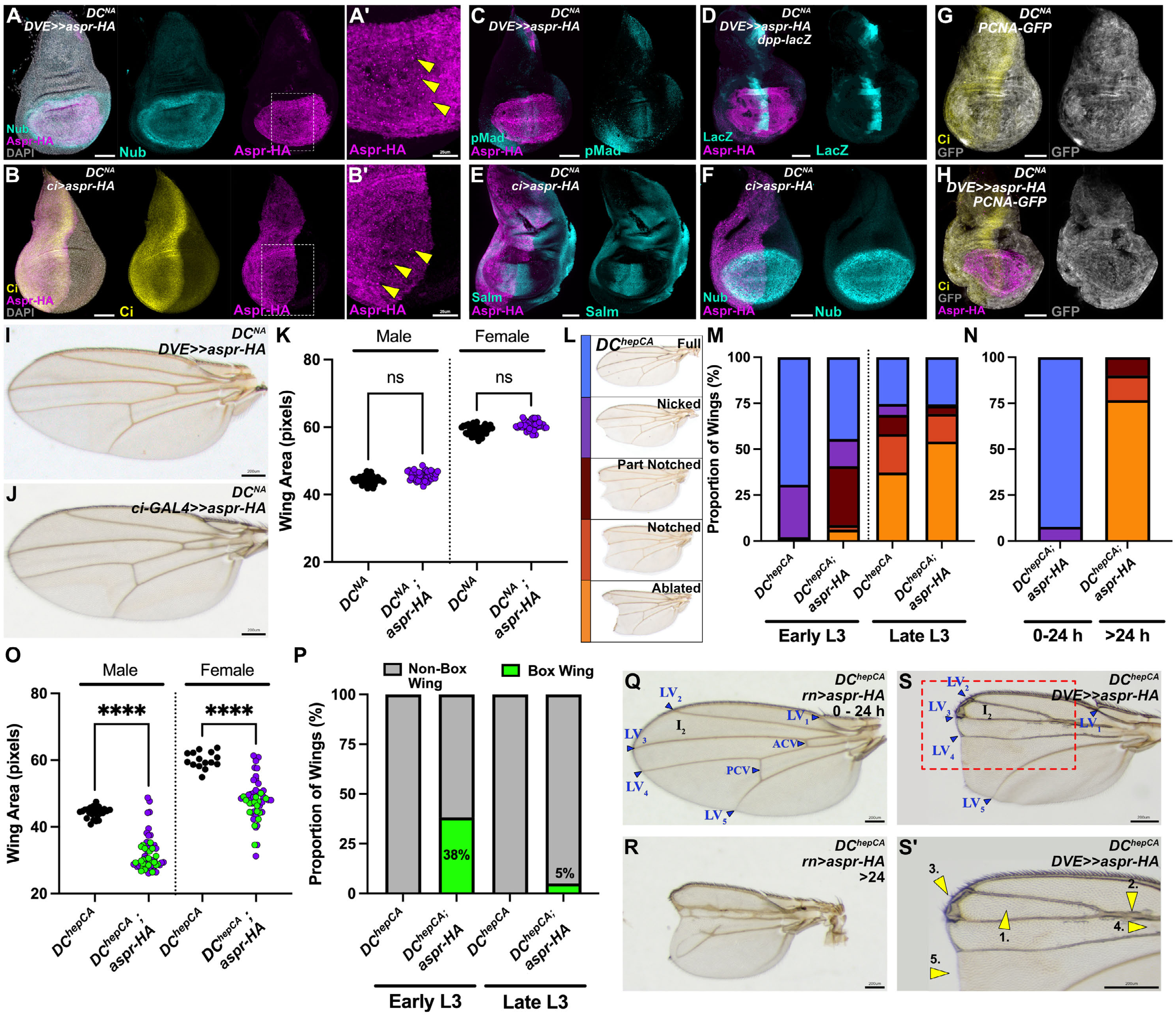
Ectopic Aspr disrupts repatterning during regeneration but not normal development. (**A–B**′) Aspr-HA (magenta) expressed with *DVE>>GAL4* (A,A′) or *ci-GAL4* (B,B′) in early L3 unablated discs (*DC^NA^*). Aspr-HA localizes to discrete puncta; higher magnification views (A′,B′) show focal accumulation (arrowheads). Co-stains: HA (magenta), Nubbin (nub, cyan, A), Ci (yellow, B); nuclei = DAPI (gray). (**C–F**) Dpp signaling (pMad and *dpp-lacZ*) and expression of *salm* and *nub* are unchanged by *aspr-HA* expression in unablated (*DC^NA^*) discs (C–F). (**G–H**) PCNA-GFP proliferation reporter is unaltered in unablated discs (*DC^NA^*) expressing *aspr-HA* with *DVE>>GAL4*. (**I–J**) Representative adult wings from undamaged discs expressing aspr-HA in the pouch (I) or anterior compartment (J) — both show normal morphology. (**K**) Quantification of wing area from experiments in (I–J); no significant change is seen with Aspr-HA (one-way ANOVA). Sample sizes: males; *y^RNAi^* n = 40, *aspr-HA* n = 44; females; *y^RNAi^*n = 41, *aspr-HA* n = 40. (**L**) Definition of regeneration scoring categories used for adult wings following ablation with DC^hepCA^. (**M**) Regeneration scoring of adult wings after ablation (*DC^hepCA^*) with pouch expression of *aspr-HA* using *DVE>>GAL4*. Aspr-HA reduces regenerative capacity in both early and late L3. Sample sizes: Early L3; *y^RNAi^*n = 50, *aspr-HA* n = 81; Late L3; *y^RNAi^* n = 67, *aspr-HA* n = 133. (**N**) Regeneration scoring of adult wings after ablation (*DC^hepCA^*) with temporally restricted *aspr-HA* expression (*rn-GAL4, GAL80ts*): 0–24 h AHS expression has no effect; expression >24 h impairs regeneration. Sample sizes: 0–24 h n = 39; >24 h n = 30. (**O**) Wing area measurements corresponding to (M); *aspr-HA* reduces wing area for both sexes and stages (one-way ANOVA; **** p < 0.0001). Sample sizes: males; *y^RNAi^* n = 25, *aspr-HA* n = 57, females; *y^RNAi^* n = 15, *aspr-HA* n = 52. (**P**) Frequency of the “box wing” phenotype in regenerated adult wings from (M): early L3 aspr-HA = 38% (box wing n = 50 of 81), late L3 aspr-HA = 5% (box wing n = 7 of 133). (**Q–R**) Representative adult wings following *aspr-HA* expression for 0-24 h (Q) or from 24 h onward (R), as in (N). Annotations indicate longitudinal veins (LV_x_) and anterior or posterior cross veins (ACV/PCV). The intervein 2 (I_2_) region is also labeled. (**S–S**′) Example box wing phenotype, red dashed outline indicates zoomed view in (S’), highlighting defects: (1) reduced I2, (2) vein fusion, (3) bristle loss, (4) missing ACV, (5) margin loss (numbered arrowheads). All scale bars = 50 μm unless noted. See also Figure S3. Full genotypes in Supplementary Genotypes.

We next examined its function during regeneration. Although endogenous *aspr* is normally downregulated after 24 h (Figures 1A–F),^31^ continuous expression of *aspr-HA* (or untagged *aspr*) throughout regeneration impaired recovery in both early and late L3 discs (Figures 3L–M, S3I–J). To determine whether this was due to the higher level of Aspr in early regeneration or its inappropriate presence in late regeneration, we used GAL80^ts^ to restrict *aspr-HA* expression to either the first 24 h AHS, or from 24 h AHS onward. Early expression had no effect (Figures 3N and Q), while late expression significantly impaired regeneration (Figures 3N and R). This suggests that while Aspr is normally required early, and ectopic Aspr at this stage has no effect, its persistence beyond 24 h disrupts recovery.

When Aspr is maintained in late regeneration, in addition to reduced regeneration, we also observed a distinct “box wing” phenotype in a subset of adults (Figures 3O–P and S–S’): ∼38% in early L3 and 5% in late L3 (Figure 3P). These wings showed consistent features that predominantly affect the anterior wing (Figures 3S-S’), including reduced intervein 2 (I_2_) between longitudinal veins 2 and 3 (LV_2_, LV_3_), 2) ectopic veins along LV_2_ and LV_3_, 3) loss of anterior wing margin bristles, 4) variable loss of ACV and PCV, and 5) a squared-off wing blade. These phenotypes also occurred with the untagged transgene in ablated wings (Figure S3K). However, these features were absent when *aspr-HA* was expressed in undamaged wings (Figures 3I–J) or when expression was restricted to early regeneration (Figure 3Q). These findings indicate that mis-patterning arises from prolonged Aspr expression. Since Aspr has only limited temporal expression during regeneration, and loss-of-function studies do not clearly reveal its specific role, we focused on the phenotypes resulting from ectopic expression to better understand the function of Aspr in regeneration.

### Aspr affects regrowth of the anterior compartment

To better understand the "box wing" phenotype resulting from *aspr* expression in late regeneration, we measured the regions of the adult wing. Posterior wing area was nearly equivalent to that of ablated wild-type controls, with only a slight reduction (Figure 4A’), whereas the anterior was significantly reduced, accounting for the majority of loss in wing size (Figure 4A). This reduction reflects smaller anterior intervein regions, most significantly of the I_2_ area (Figure 4B–C). I_3_ and I_4_ were only measured when the ACV was present. To confirm that Aspr affects anterior patterning and growth, we drove *aspr-HA* solely in the anterior or posterior compartment using *ci-GAL4* or *hh-GAL4*. Anterior-specific expression during regeneration produced phenotypes resembling those of whole-pouch expression: I_2_ reduction, LV_2_–LV_3_ fusion and thickening, ectopic vein tissue, ACV loss, and absent anterior margin bristles (Figure 4D). Posterior-specific expression caused only a small reduction in the I_6_ area (Figure 4E), matching the mild posterior reduction observed in whole-pouch expression (Figure 4A’), but shows no other patterning defects. Expression with either driver in undamaged discs caused no wing phenotypes (Figure S3G and S3H). Thus, the overall box wing phenotype likely reflects a combination of the significant anterior and mild posterior effects (Figure 4D and 4E). Together, these findings strongly suggest that maintaining *aspr* specifically disrupts anterior growth and patterning during regeneration.

**Figure 4.**
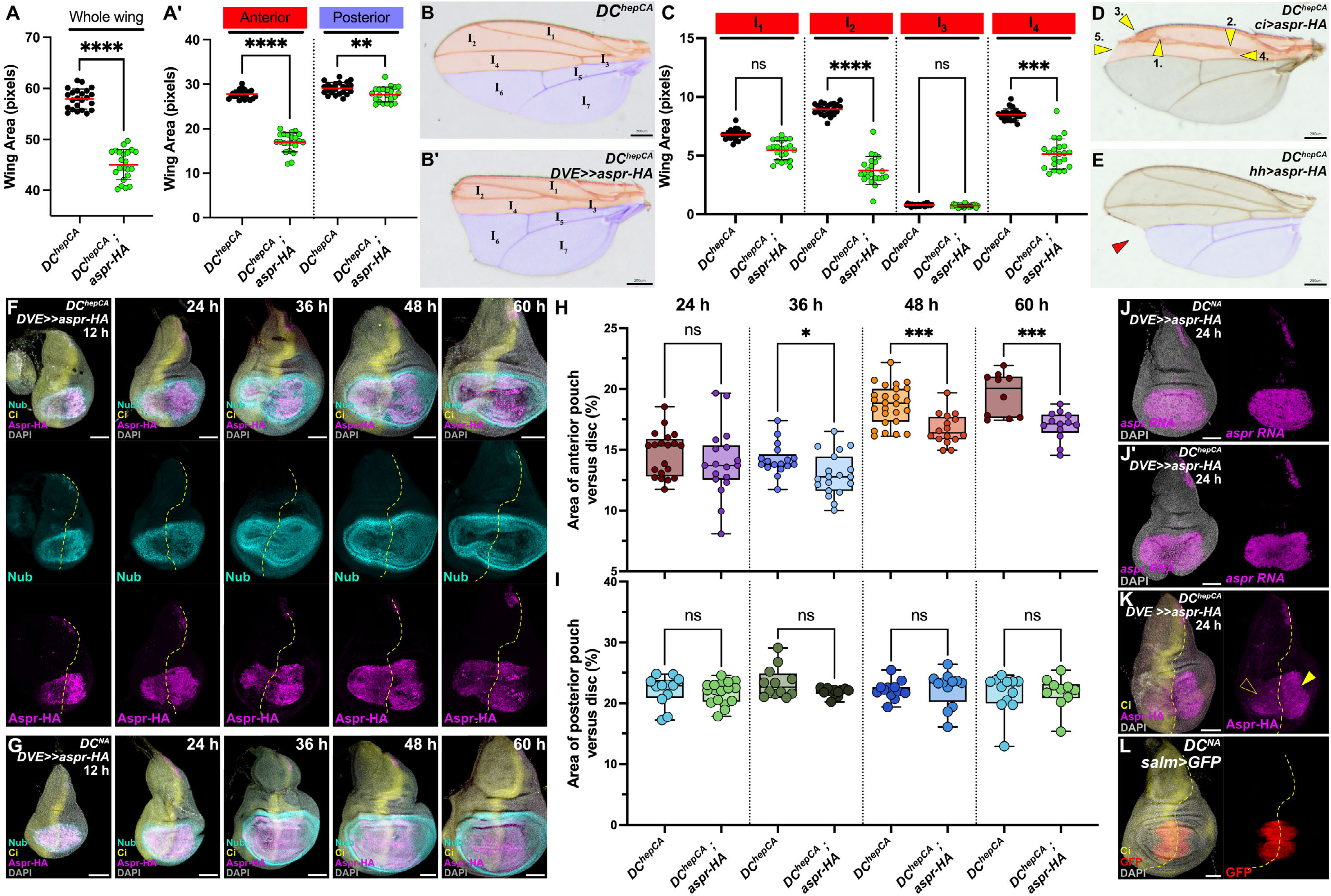
Ectopic Aspr has an anterior-specific effect on regeneration. (**A–A**′) Wing area measurements of adult female wings from early L3 ablated discs (*DC^hepCA^, DVE>>GAL4*): (A) box-wing phenotype (green) with Aspr-HA is significantly smaller than control wings (black); (A′) anterior and posterior areas measured separately show a larger reduction in the anterior. Statistics: one-way ANOVA. Sample sizes: *y^RNAi^* n = 23, *aspr-HA* n = 22. **** p < 0.0001; anterior **** p < 0.0001; posterior ** p = 0.0040. (**B–B**′) Representative adult wings from ablated discs (*DC^hepCA^, DVE>>GAL4*) expressing *y^RNAi^* (B) or *aspr-HA* (B′). Anterior (red) and posterior (blue) compartments shaded. Intervein regions labeled. (**C**) Quantification of anterior intervein regions I_1_–I_4_ from (A′). I_2_ and I_4_ are significantly reduced in *aspr-HA* wings (one-way ANOVA). Sample sizes: I_1_; y^RNAi^ n = 15, *aspr-HA* n = 12; I_2_; *y^RNAi^* n = 20, *aspr-HA* n = 18; I_3_; *y^RNAi^* n = 17, *aspr-HA* n = 19; I_4_; *y^RNAi^* n = 26, *aspr-HA* n = 16. I_1_, I_3_ = ns; I_2_ = **** p = 0.001; I4 = *** p = 0.001. (**D–E**) Anterior (*ci-GAL4*) *aspr-HA* expression after ablation produces multiple anterior patterning defects (numbered arrowheads) seen in box wings; posterior (*hh-GAL4*) *aspr-HA* only reduces wing area, without the anterior patterning defects (red arrowhead). (**F–G**) Regenerating disc time course (*DC^hepCA^, DVE>>GAL4*) stained for HA (magenta), Nub (cyan), and Ci (yellow). Anterior pouch is consistently smaller from 12–60 h AHS with Aspr-HA (F), whereas developmental control (*DC^NA^, DVE>>GAL4*) shows no change across time points with Aspr-HA (G). A/P boundary indicated by Ci. (**H–I**) Quantification of pouch compartments (anterior as % total disc in H; posterior in I) from the time course in (F) versus control. Anterior pouch is significantly reduced at 36 h, 48 h, and 60 h AHS; posterior pouch is unchanged. Sample sizes per timepoint: 24 h; *y^RNAi^*n = 20, *aspr-HA* n = 18; 36 h; *y^RNAi^* n = 17, *aspr-HA* n = 19; 48 h; *y^RNAi^* n = 26, *aspr-HA* n = 16; 60 h; *y^RNAi^* n = 10, *aspr-HA* n = 13. Anterior: 24 h = ns, 36 h * p = 0.0337, 48 h *** p = 0.0005, 60 h *** p = 0.0007. (**J–K**) *aspr* RNA (magenta) at 24 h AHS in unablated (*DC^NA^*) and ablated (*DC^hepCA^*) discs expressing *aspr-HA*; transcript levels appear similar between conditions (J). Aspr-HA protein (K) is reduced in the anterior (open arrowhead) relative to posterior (arrowhead); Ci outlines A/P boundary. (**L**) Unablated disc with R85E08>GFP (red) shows transgene expression from the ablation domain spans both compartments similarly; Ci marks the A/P boundary. All scale bars = 50 μm unless otherwise noted. See also Figure S4. Full genotypes in Supplementary Genotypes.

To examine the origin of these phenotypes, we monitored regeneration over time with *aspr-HA* expression and stained for *nub*, *ci*, and HA to mark the pouch, anterior compartment, and Aspr, respectively (Figure 4F). The anterior pouch is consistently undergrown from 12–60 h AHS, while the posterior regenerated similarly to controls. Quantification confirmed this observation, with significant anterior undergrowth at all time points except 24 h (Figure 4H and 4I). This aligns with the idea that persistent Aspr only disrupts late regeneration, when endogenous *aspr* is normally downregulated. Ectopic *aspr* without damage has no effect on pouch size (Figures 4G, S4A–B), consistent with the absence of adult phenotypes (Figure 3I). To explore this anterior-specific effect, we examined Aspr localization. Although *aspr-HA* transcripts are equally expressed in both compartments (Figure 4J–J’), Aspr-HA protein appeared weaker in the anterior pouch of ablated discs (Figure 4K), which could indicate compartment-specific protein localization, stability or usage. We ruled out differences in ablation efficiency, as the *salm* enhancer used to induce damage drives similarly in both compartments (Figure 4L). These results indicate that maintaining Aspr beyond its normal early timing limits regrowth specifically in the anterior pouch. While the reason for this compartment-specificity remains unclear, these data suggest that the normal role of Aspr is to support early regenerative growth in the wing pouch.

### Aspr functions to regulate regeneration-specific pouch growth through *vestigial*

To better understand the growth phenotype associated with Aspr during regeneration, we used an E2F reporter (*PCNA-GFP*) to monitor proliferation over time, with Ci staining to mark compartment boundaries. In control discs, injury triggers strong proliferation near the wound, indicative of a regeneration blastema (Figure 5A). This proliferation peaks over 24-48 h and returns to surrounding levels by 60 h (Figures 5A–A’’’ and C). We noted that proliferation seems biased toward the anterior, consistent with previous reports showing differences in proliferative potential between compartments.^35–37^ Ectopic *aspr-HA* expression reduces proliferation in both compartments, particularly in the anterior (Figure 5B–B’’’ and C), matching the undergrowth seen in discs and adult wings (Figures 4F and 4H). *aspr-HA* had no effect on proliferation in undamaged discs (Figures 3G and 3H). We attempted to repeat these experiments with anti-PH3 and EdU incorporation, but neither was able to clearly demonstrate these differences (Figures S5A–L), likely because the E2F reporter integrates proliferative activity over time, while PH3 and EdU capture only transient mitotic or S-phase events, making subtle differences harder to detect.

**Figure 5.**
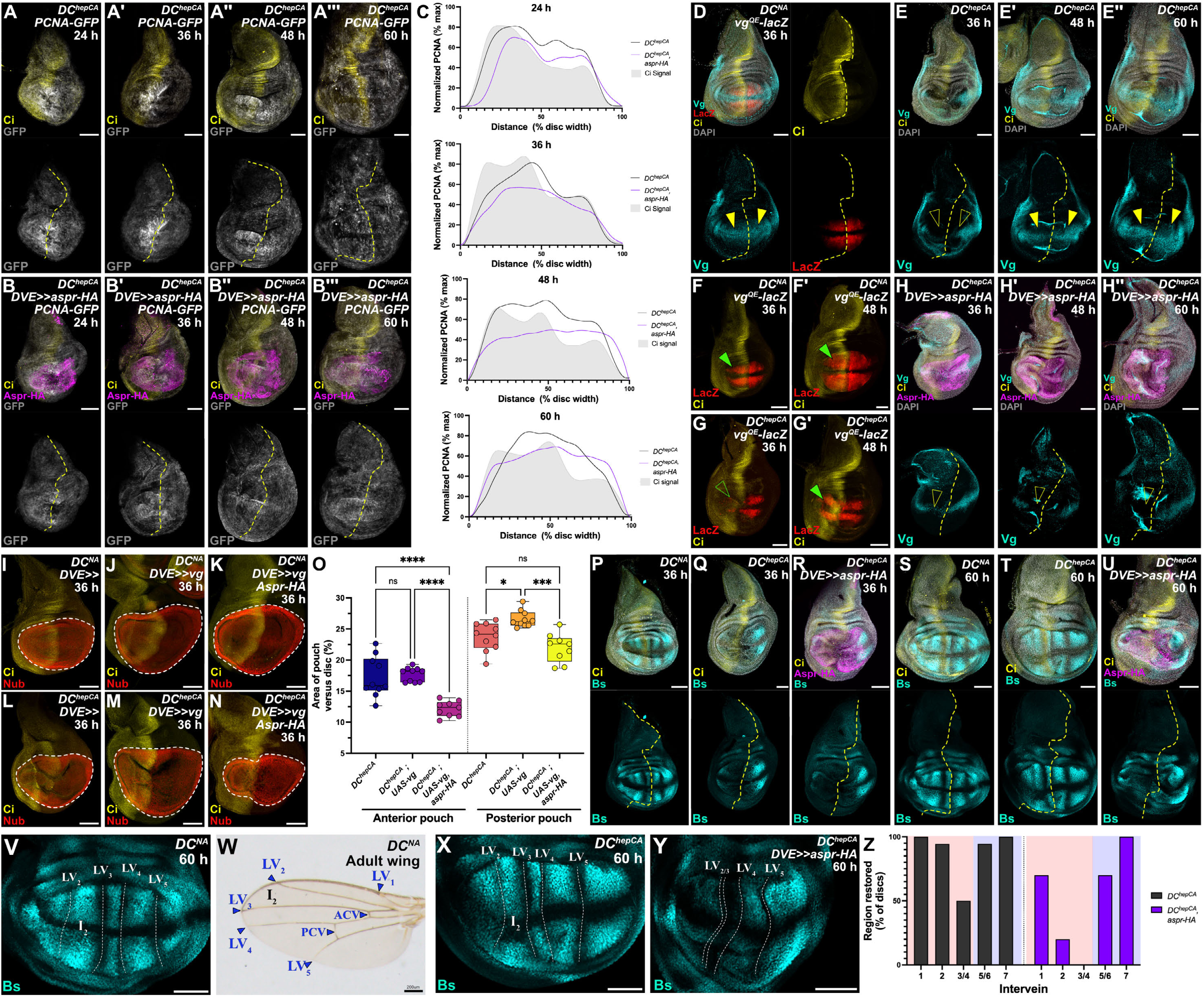
Aspr impairs anterior proliferation and Vestigial restoration during regeneration. (**A–B**L) Time course of PCNA-GFP in early L3 ablated discs (*DC^hepCA^, DVE>>GAL4*): controls (A–A_) show elevated proliferation at 24–48 h with slight anterior bias that declines by 60 h; *aspr-HA* expressing discs (B–B_) show reduced PCNA-GFP fluorescence and loss of anterior bias. A/P boundary = Ci (yellow dotted line). (**C**) LOWESS splines of PCNA-GFP intensity across disc width for each timepoint (n = 4 discs/timepoint) illustrate loss of anterior bias in aspr-HA discs. (**D–G**′) *vg^QE^-lacZ* and *vg* expression dynamics: unablated discs (*DC^NA^,* D and F) show symmetric Vg and lacZ across time points (arrowheads); ablated discs (*DC^hepCA^*, E-E’’ and G-G’) lose Vg at 36 h (E), which is restored by 48–60 h in controls (E′-E’’). *vg^QE^-lacZ* is lost in the anterior at 36 h (G, open arrowhead), restored by 48 h (G’, arrowhead). (**H–H**′′) Ablated discs expressing *aspr-HA* (magenta, *DC^hepCA^, DVE>>GAL4*) show persistent loss of anterior Vg (cyan) from 36–60 h post-ablation (open arrowheads). Ci (yellow) indicates anterior. (**I–N**) Unablated (*DC^NA^, DVE>>GAL4*, I–K) and ablated (*DC^hepCA^, DVE>>GAL4*, L–N) discs at 36 h AHS expressing *vg* or *vg+aspr-HA*, stained for Ci (yellow) and Nub (red). Vg induces pouch overgrowth in unablated discs, which is limited by co-expression of *aspr-HA* in ablated discs, particularly in the anterior. (**O**) Quantification of pouch compartments for the conditions in (I–N). Anterior and posterior pouch areas (as % of total disc) are reduced by Aspr-HA, with a larger effect in the anterior. Sample sizes: anterior; *y^RNAi^* n = 10, *vg* n = 10, *vg+aspr-HA* n = 9 ; posterior; *y^RNAi^* n = 10, *vg* n = 10, *vg+aspr-HA* n = 9. Statistics: one-way ANOVA. Annotated p-values: anterior **** p < 0.0001; posterior * p = 0.0373, *** p = 0.0002, **** p < 0.0001; ns = not significant. (**P–U**) Blistered (Bs, cyan) marks intervein regions in unablated discs (*DC^NA^*, P and S), and ablated discs (*DC^hepCA^, DVE>>GAL4)* at 36 h (Q and R) or 60 h (T and U) AHS. Following ablation, Bs is lost at 36 h (Q) and restored by 60 h in control discs (T) but remains absent in *aspr-HA* expressing discs (U). Ci (yellow) indicates anterior. (**V**) High magnification view of unablated disc (*DC^NA^*) at 60 h with I_2_ intervein, longitudinal veins (LVx) labelled, indicated by Bs staining (cyan). (**W**) Adult wing with labels corresponding to disc in (V), and anterior/posterior cross veins (A/P CV). (**X-Y**) High magnification view of ablated discs (*DC^hepCA^, DVE>>GAL4*) at 60 h with I_2_ intervein, and longitudinal veins (LVx) labelled, indicated by Bs staining (cyan). Control discs restore anterior Bs and I2 (X), while those expressing *aspr-HA* discs fail to restore Bs or reestablish I_2_ (Y). (**Z**) Quantification of intervein restoration across I_1_–I_7_: controls (black) show robust restoration; Aspr-HA (magenta) reduces restoration especially in anterior I_2_ and I_3/4_. All scale bars = 50 μm unless otherwise indicated. See also Figure S5. Full genotypes in Supplementary Genotypes.

We next screened candidate regulators of growth that might participate in regeneration, and found that *vestigial (vg)*, a key wing selector gene, is disrupted upon *aspr* misexpression. *vg* is normally expressed along the D/V boundary (Figure 5D), and is patterned by *wg*, *notch*, *dpp*, and autoregulation.^38–41^ After DC^hepCA^ ablation, Vg is reduced until 36 h AHS but restored by 48 h (Figures 5E–E’’). However, with sustained *aspr-HA* expression, Vg fails to reestablish in the anterior pouch through 60 h AHS and often remains absent until pupariation (Figures 5H–H’’). Interestingly, the *vg* quadrant enhancer (vg^QE^), which initiates *vg* expression in response to Wg,^38^ also shows anterior-specific loss after damage, typically recovering by 48 h AHS in controls (Figures 5F–G’ and S5M-T). Due to genetic constraints, we could not assess vg^QE^ reporter activity in *aspr-HA*-expressing discs, but these results suggest that anterior *vg* is less robustly reinitiated after injury, and thus may be more sensitive to prolonged Aspr expression. To test whether Aspr can suppress Vg-mediated growth, we expressed *vg* in the presence and absence of *aspr-HA* in damaged and undamaged discs, and quantified pouch size. Ectopic expression of *vg* alone yields larger pouch size in both undamaged (Figure 5I–J and S5U), and damaged discs (Figure 5L–M and 5O). However, co-expression with *aspr-HA* significantly suppresses this growth in damaged discs (Figure 5N–O), which does not occur in undamaged discs (Figure 5K and S5U). This growth suppression primarily occurs in the anterior compartment (Figure 5N–O), consistent with the previous observations of compartment-specific differences in Vg restoration (Figure 5E–H’’). Together, these data confirm that 1) Aspr regulates Vg-mediated growth in the pouch, 2) this effect occurs preferentially in the disc anterior, and 3) that the function of Aspr is most significant in damaged discs that are undergoing regeneration. However, whether Aspr modulates Vg itself, or rather influences an intermediate factor that participates in Vg-dependent growth, is unclear (see later).

Although anterior pouch tissue is reduced with ectopic Aspr, it still acquires vein/intervein identity (Figures 3S-S’). To assess patterning, we examined *blistered (bs/DSRF)*, which represses vein fate to define intervein regions (Figure 5V–W).^42,43^ In undamaged discs, the vein/intervein regions are defined by *bs* expression at 36 h AHS and is unchanged at 60 h AHS (Figures 5P, S and V). After damage, *bs* is transiently lost but returns by 48-60 h AHS (Figures 5Q, T and X). *aspr-HA* expression has no clear effect through 36 h (Figure 5R), but by 60 h, I_2_ fails to form and LV_2_ and LV_3_ are fused (Figures 5U and Y), mirroring adult wing defects. Since most of the pouch retains proper *bs* expression and vein identities despite overlapping Aspr (Figures 5U, Y and Z), this patterning defect possibly results indirectly from impaired growth. Alternatively, as *vg* has been shown to regulate *bs,*^39^ the phenotype may stem from disrupted *vg* expression. Together, these findings suggest that Aspr normally promotes proportional growth during early regeneration, partly by supporting timely *vg* reactivation, and when mis-regulated it impairs both regenerative growth and patterning.

### Aspr limits expression of late-acting differentiation-associated genes specifically during regeneration

Changes in Vg may account for anterior pouch size reduction and vein/intervein patterning defects in the box wing phenotype, but significant margin abnormalities and loss of bristles is also observed (Figures 3S–S’). To investigate this, we examined genes involved in margin cell differentiation. Bristles at the wing margin derive from sensory organ precursors specified by Notch and Wg signaling.^44^ Previously, we showed that Aspr may affect expression timing of the Notch and Wg target gene *cut* (*ct*) at the margin specifically following injury (Figure 6A).^22^ We revisited this by testing *aspr-HA* expression during regeneration. Normally, Ct is lost following ablation and restored in all discs by 60 h AHS (Figures 6B–C). With *aspr-HA* expression, Ct is fully restored in only 18% of discs (n = 50), with most showing only partial recovery (Figure 6D). *aspr-HA* had no effect on Ct in undamaged discs (Figure S6A), confirming it interferes with *ct* reactivation during regeneration. Furthermore, restricting expression of *aspr-HA* solely to the *ct* domain using *ct-GAL4* results only in the loss of bristles component of the box wing phenotype (Figure S6F), which we attribute to Aspr’s repressive effect on Ct. This phenotype is not present when *aspr-HA* is driven in the *ct* domain in undamaged discs (Figure S6E). *ct* is maintained indirectly by Wg via the Notch ligands Delta and Serrate (Figure S6H).^45^ Delta expression, normally restored by 60 h AHS (Figure 6E–F), is also suppressed by *aspr-HA* (Figure 6G), which may explain the persistent loss of Ct. As both *vg* and *Delta* (and thus indirectly *ct*) are targets of Wg, we examined other differentiation-associated Wg-responsive genes. *aristaless* (*al*) encodes a homeobox transcription factor downstream of Wg^46^. al failed to be restored with *aspr-HA* by 60 h AHS (Figure 6L–N). Similarly, *achaete* (*ac*), a Wg-responsive factor that promotes neural fate^47,48^ normally appears after 48 h time point in undamaged discs (Figure S6I), is reestablished in wild type ablated discs by 60 h AHS (83.9% of discs, Figure 6H–J), but fails to be reestablished in most *aspr-HA*-expressing discs at this time point (7.9%) (Figure 6K). This finding, alongside the reduction in Ct (Figure 6D) likely underlies the reduced bristle counts observed in wings (Figure S6G). Thus, prolonged *aspr* disrupts the reactivation of multiple differentiation-promoting genes (Figure 6O), all of which are directly or indirectly downstream of Wg signaling. None of these genes were altered by *aspr-HA* expression in undamaged discs (Figures S6A-S6D). Many Wg targets also depend on Dpp, however, expression of a *Dpp-lacZ* reporter, pMad, and its target *spalt* (*salm*) are unaffected by *aspr-HA* during development or regeneration (Figures 6L–N and S6J–O’’). Given the strong overlap between *wg* and *aspr* expression in the blastema,^31^ and the fact that several affected genes lie downstream of Wg, we hypothesize that Aspr normally delays Wg-driven differentiation and sustains early regenerative growth through regulation of Vg.

**Figure 6.**
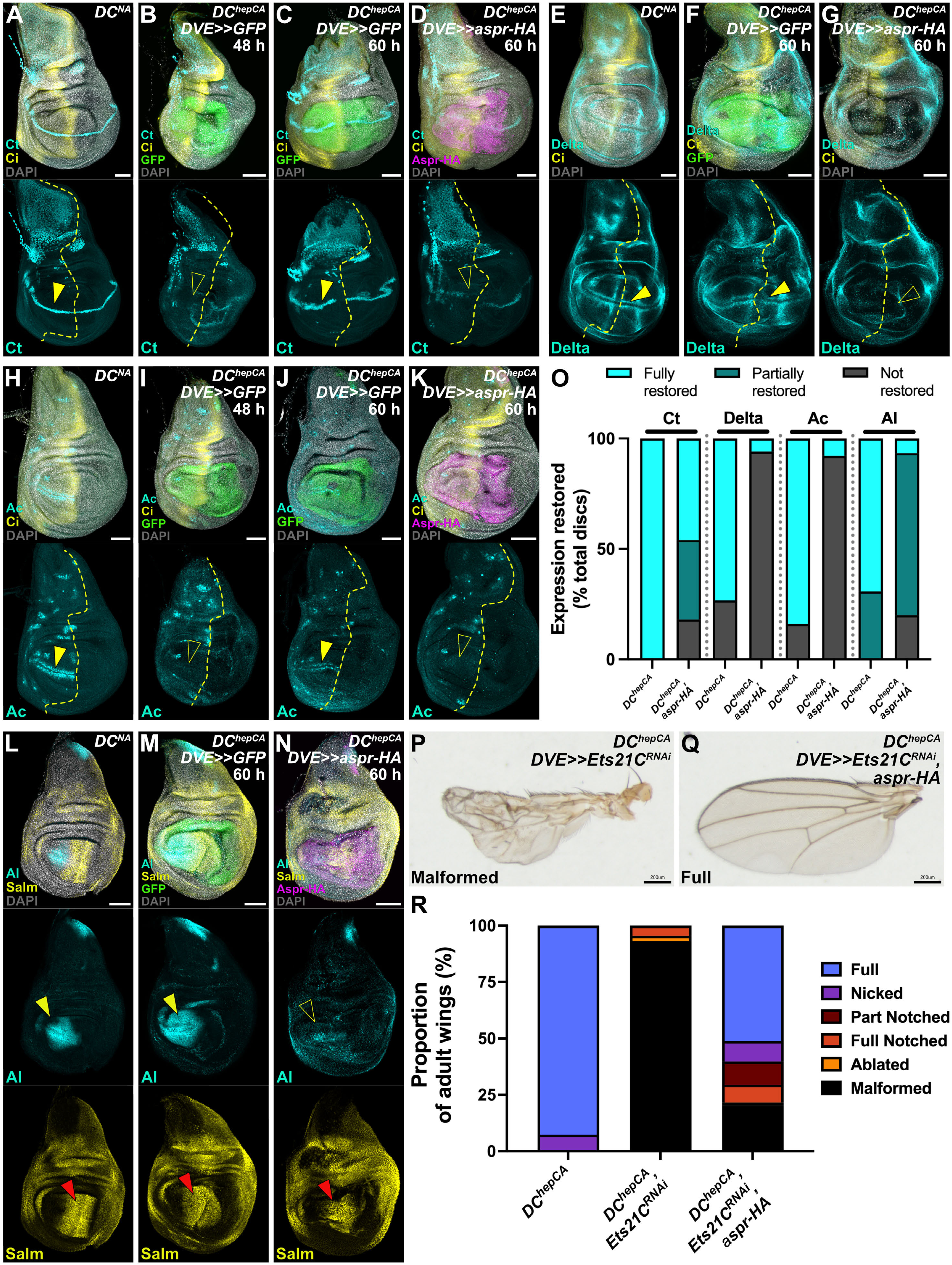
Aspr inhibits restoration of anterior cell-fate markers during regeneration. (**A**) Unablated (*DC^NA^*) disc stained for Cut (Ct, cyan) showing margin expression including anterior (arrowhead). Ci (yellow, dotted line) marks the A/P boundary; nuclei = DAPI (gray). (**B–D**) Ablated discs (*DC^hepCA^, DVE>>GAL4*) at 48 h (B) and 60 h (C,D) AHS stained for Ct. In GFP controls, anterior Ct is absent at 48 h (B, open arrowhead) and restored at 60 h (C, arrowhead). Ectopic Aspr-HA blocks Ct restoration at 60 h (D, open arrowhead). (**E–G**) Delta (Dl) staining: unablated (E) shows margin/vein expression; at 60 h post-ablation Dl is restored in controls (F, arrowhead) but not in aspr-HA discs (G, open arrowhead). (**H–K**) Discs as in (B–D) showing Achaete (Ac) dynamics: Ac is absent at 48 h post-ablation (I, open arrowhead) and restored by 60 h in controls (J, arrowhead); Aspr-HA prevents restoration at 60 h (K, open arrowhead). (**L–N**) Aristaless (Al) and Salm: Al is anterior-restricted and restored by 60 h in controls (M, yellow arrowhead); aspr-HA blocks Al restoration (N, open arrowhead) but does not affect Salm (N, red arrowhead). (**O**) Quantification of marker restoration at 60 h for Ct, Dl, Ac, and Al. Y-axis indicates percent discs with full (cyan), partial (turquoise), or no restoration (dark gray). Ectopic Aspr-HA reduces restoration of all anterior markers. Sample sizes: Ct: *y^RNAi^* (n = 28), *aspr-HA* – Full (n = 23), Partial (n = 18), None (n = 9) Dl: *y^RNAi^* – Full (n = 11), None (n = 4); *aspr-HA* – Full (n = 1), None (n = 16) Ac: *y^RNAi^* – Full (n = 47), None (n = 9); *aspr-HA* – Full (n = 3), None (n = 35) Al: *y^RNAi^* – Full (n = 9), Partial (n = 4); *aspr-HA* – Full (n = 1), Partial (n = 11), None (n = 3) (**P–Q**) Representative adult wings after early L3 ablation (*DC^hepCA^*) with *Ets21C*^RNAi^ alone (P) or combined with *aspr-HA* (Q). *Ets21C* knockdown impairs regeneration; co-expression of *aspr-HA* rescues malformed wings. (**R**) Regeneration scoring corresponding to (P–Q). Sample sizes: Control n = 27, *Ets21C*^RNAi^ n = 44, *Ets21C*^RNAi^ + *aspr-HA* n = 88. All scale bars = 50 μm unless otherwise indicated. See also Figure S6. Full genotypes in Supplementary Genotypes.

The idea that Aspr acts to delay differentiation is supported by previous work showing that *aspr* is downstream of the ETS transcription factor Ets21C, which promotes regeneration by supporting the expression of genes like *Mmp1*, *upd3*, *pvf1*, and *aspr*, while preventing premature induction of repatterning genes like *rotund* and *nubbin*.^31^ As with *aspr*, mutation of *Ets21C* causes defective, undersized wings post-ablation, but does not affect development. Knockdown of *Ets21C* in ablated wings shows this phenotype (Figures 6P and R), but this is almost entirely rescued by co-expression of *aspr-HA* (Figures 6Q and R), suggesting that a key function of Ets21C in regeneration is the activation of *aspr* to prevent premature differentiation and promote tissue repair.

### Aspr protein structure suggests a role in protein trafficking and secretion

Aspr appears to regulate growth and differentiation during regeneration, but its molecular function remains unclear. To investigate this, we analyzed its predicted protein structure using AlphaFold (Figure 7A and S7A), which reveals a solenoid-like fold with multiple epidermal growth factor (EGF)-like domains (Figure 7B). InterProScan and SMART predict seven EGF domains, while Prosite identifies four. These domains are typical of secreted or membrane-associated proteins involved in signaling, adhesion, and matrix organization^49^, supporting a potential extracellular role for Aspr. To further test this, we used six localization prediction tools. Four of these (SignalP, Phobius, Protter, Deeploc) predict a cleavable N-terminal signal peptide with a high-confidence cleavage site between Gly23 and Lys24 (Figures 7C and S7B–E). TargetP and TMHMM also supported this with moderate confidence (∼0.5) (Figures 7C and S7F–G). These tools also consistently predict Aspr to be extracellular (Figure 7C) reinforcing the idea it is likely secreted. Homology searches identify several structurally related extracellular proteins: human NELL2, human/mouse EDIL3, and *Drosophila* Dumpy (Figure 7D). Alignments of their signal peptides with that of Aspr revealed shared biochemical properties and α-helical structures typical of signal sequences (Figure 7D and S7H–L), while RMSD comparisons showed strong similarity to Dumpy (0.189), NELL2 (0.248), mouse EDIL3 (0.635), and human EDIL3 (1.04) (Figure 7E), supporting the idea that Aspr has structural conservation with other secreted proteins. Together, these structural analyses are consistent with Aspr functioning as a secreted or membrane-associated extracellular protein.

**Figure 7.**
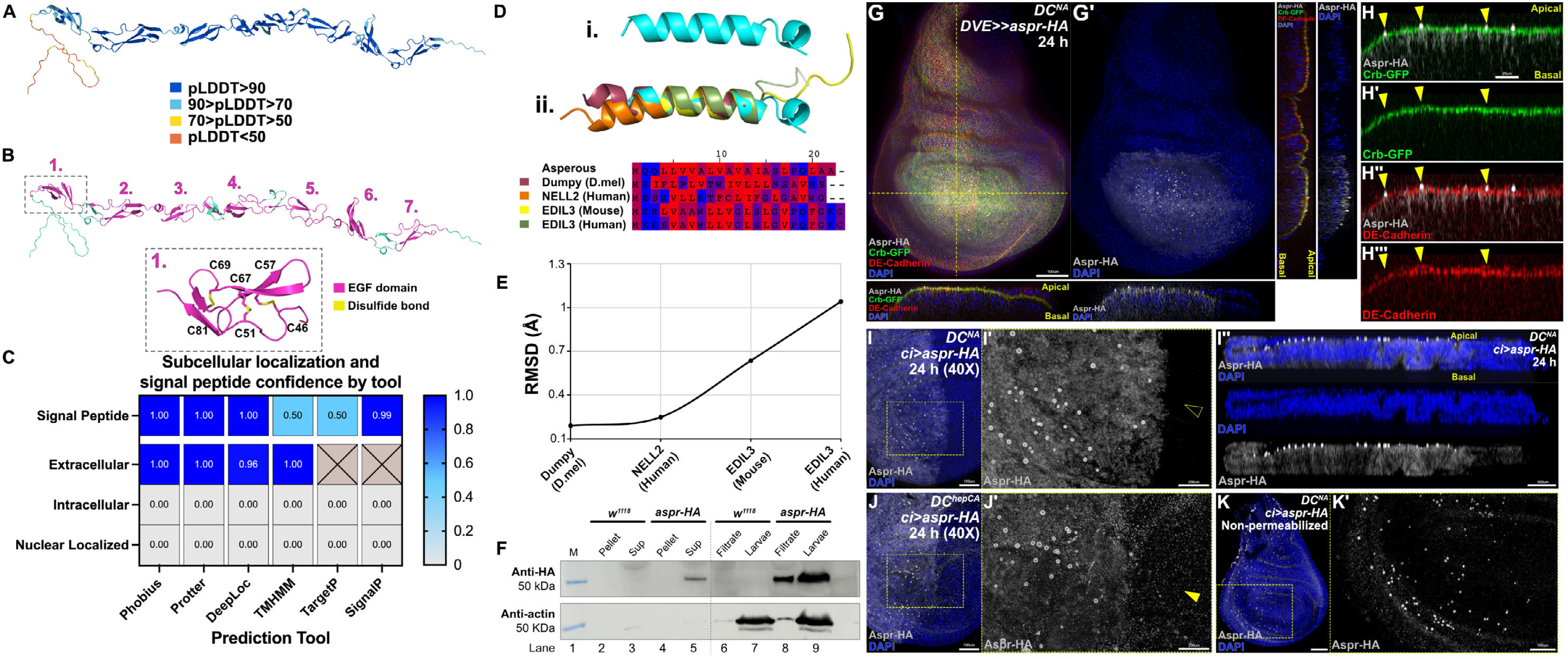
Structural and in vivo analyses show Aspr is an extracellular protein. (**A**) AlphaFold predicted structure of full-length Aspr colored by pLDDT confidence: very high (>90, blue), confident (70–90, cyan), low (50–70, yellow), very low (<50, orange). (**B**) AlphaFold model highlighting seven predicted EGF-like domains (magenta) separated by non-EGF regions (cyan). Inset shows EGF domain 1 with predicted disulfide bonds (yellow) and cysteines. (**C**) Heatmap summarizing signal peptide and subcellular localization predictions from six tools (SignalP, Phobius, TargetP, Protter, TMHMM, DeepLoc); scores range 0 (low, blue) to 1 (high, yellow); grey = no data. **D**) (i) Structural alignments of the Aspr signal peptide (cyan) with signal peptides from Dumpy isoform V (*Drosophila*, maroon), NELL2 (human, orange), and EDIL3 (mouse, yellow; human, green). (ii) Sequence alignment showing conserved features (e.g., polarity, hydrophobicity), with residues colored by chemical class: hydrophobic (dark red), mildly hydrophobic (red), polar (blue), moderately polar (purple). (**E**) RMSD values from structural alignments in (D). Each point represents structural similarity between Aspr and comparison proteins; lower values indicate greater similarity. (**F**) Western blot of hemolymph fractions from control (*w^1118^*) and *ci-GAL4>aspr-HA* larvae, probed with HA and Actin antibodies. HA is detected in whole *aspr-HA* larvae (lane 9) and filtered hemolymph (lane 8), and in the supernatant (lane 5) but not the pellet (lane 4) of hemolymph after ultracentrifugation, consistent with secretion. No HA is detected in control larvae (lanes 2–3, 6–7). Actin is detected in whole larvae and filtrate (lanes 8–9). (**G–G’**) Unablated (*DC^NA^, DVE>>GAL4*) discs with the Crb-GFP apical marker (green), expressing *aspr-HA* (grey). DE-Cadherin labels cell junctions (red); DAPI (blue) labels nuclei. (G) Yellow dotted line indicates imaging planes shown in cross sections. Aspr-associated EVs are found apical to Crb-GFP, which is apical to DE-Cadherin. (**H–H’’’**) High magnification of transverse sections in (G). Aspr-associated EVs (yellow arrowheads) occur apically to the Crb marker (H-H’) and DE-Cadherin (H’’-H’’’). Aspr also appears apically within disc cells. (**I–I’’**) High magnification imaging of unablated (DC^NA^) disc expressing *aspr-HA* at 24 h via ci-GAL4. (I) Surface view shows vesicles; yellow box highlights zoomed region in (I’). Open arrowhead marks area lacking vesicles beyond expression domain. (I’’) Cross-section reveals vesicles localize to the apical surface. (**J-J’’**) High magnification imaging of ablated (*DC^hepCA^*) disc expressing *aspr-HA* at 24 h via ci-GAL4. (J) Surface view shows increased vesicles, with zoomed region in (J’). Arrowhead indicates small vesicles outside the expression domain. (**K-K’**) Genotype as (I) imaged without permeabilization. Aspr-HA vesicles are visible (zoomed in J’) while the more diffuse intracellular cytoplasmic signal is lost, consistent with Aspr having an extracellular localization. All scale bars represent 50 μm unless otherwise indicated. See also Figure S7. Full genotypes are provided in the Supplementary Genotypes file.

We experimentally tested Aspr secretion using western blots on hemolymph bled from larvae expressing *aspr-HA* (*ci-GAL4>UAS-aspr-HA*). HA was detected in both filtered hemolymph and larval debris (Figure 7F, lanes 8-9), confirming previously published proteomic analysis that identified Aspr in hemolymph^50^. We further processed the filtered hemolymph by subjecting it to additional high-speed centrifugation, which separates extracellular proteins and structures like vesicles away from cells and larger debris. Repeating HA detection on this sample shows that Aspr is present in the supernatant (Figure 7F, lane 4-5), confirming its extracellular nature. Control larvae (*w^1118^*) lacked HA signal (Figure 7F, lanes 2-3, 6-7), while anti-Actin was used to indicate cellular presence in each sample (trace Actin in filtrates likely reflects cells such as hemocytes). Together, these findings strongly support Aspr as a secreted extracellular protein.

To validate the localization of Aspr *in vivo*, we re-examined its distribution in the wing disc, focusing on the punctate structures previously observed (Figures 3A–B). High-resolution imaging of undamaged discs expressing *aspr-HA* revealed spherical extracellular vesicle (EV)-like structures (300–1400 nm diameter, Figure S7M–P) located above the apical surface (Figure 7G and I–I’’), confirmed by co-staining with the apical marker Crumbs (*crb-GFP*) in transverse sections (Figure 7H–H’’’) and a membrane-associate GFP reporter (*CD8::GFP*, Figure S7Q–R’’’). Notably, EVs do not incorporate this GFP label (Figure S7R–R’), indicating that they are unlikely to arise by simple outward budding from the labeled apical plasma membrane, but instead may originate from intracellular compartments via a secretory route, such as exosome-like vesicle release.^51^ Although our anti-Aspr antibody has limited sensitivity, we successfully detected EVs using this antibody in *aspr-HA*-expressing discs (Figure S8A-S8B’), ruling out artifacts of HA staining. This was further validated using different anti-HA antibodies and control staining conditions (Figure S8C and S8C’). Most importantly, extracellular vesicles are also detected using the anti-Aspr antibody in ablated discs expressing only endogenous *aspr*, without ectopic *aspr-HA* (Figure S8D–S8D″). These EVs are smaller (typically <100 nm in diameter), more uniform in size, and less frequent, exhibiting features of exosome-sized vesicles rather than larger plasma membrane–derived microvesicles,^26,29^ and suggesting that ectopic *aspr-HA* expression increases both EV size and abundance. These observations support the presence of Aspr-containing EVs under physiological damage conditions, but additional markers and functional assays will be required to determine their precise biogenesis, secretion route, and role in regeneration.

Aspr-associated EVs appear alongside intracellular Aspr, which is enriched apically in disc cells (Figure 7G–H’’). An extracellular-optimized staining protocol using non-permeabilized discs eliminates the intracellular Aspr signal, allowing clearer visualization of EVs outside the cells (Figure 7K-K’), located between the disc proper and the overlying peripodial membrane (Figures 7I–I’’). These EVs form in the pouch, hinge, and notum when *aspr-HA* is expressed throughout the anterior compartment via *ci-GAL4*, but are most frequently observed in the pouch (Figures 7I–I’ and 3B-B’), and can also be detected in other imaginal discs like the leg (Figure S8E). Notably, EVs are present in both undamaged and regenerating discs expressing *aspr-HA* (Figure 7I-J’). However, regenerating discs show a significantly greater number of the smaller EVs, including in regions lacking active *aspr-HA* expression (Figure 7J-J’), consistent with the possibility that extracellular Aspr may be able to disperse more broadly beyond its site of expression in regenerating discs. These EVs are distinct from the cellular debris generated by ablation (Figure S8H–J’). Overall, these structural, computational, and biochemical results demonstrate that Aspr is a secreted protein, and that it forms EVs in the wing disc, particularly during regeneration, consistent with its proposed role in extracellular signaling.

### Aspr co-localizes with Wg and regulates WNT signaling

Maintaining *aspr* expression into late regeneration affects growth and patterning in ways consistent with altered Wg signaling (Figures 5 and 6). Given the predicted protein structure and localization of Aspr to secreted EVs (Figure 7), we tested whether Aspr influences Wg signaling via an extracellular mechanism. Wg is secreted and signals through tightly regulated processes involving chaperones,^52^ lipoprotein particles,^53^ cytonemes^54,55^ juxtracrine mechanisms,^56^ and via EVs such as exosomes.^26,29,57,58^ To examine Wg regulation by Aspr, we expressed *aspr-HA* during development and regeneration while staining for Wg. In undamaged discs, *aspr-HA* has no effect on Wg levels (Figures S9A–C). In regenerating discs, however, Wg protein appears elevated from 18-48 h AHS, especially in the anterior pouch (Figures 8A–D). This is despite *aspr-HA* being driven throughout the pouch. By 60 h AHS, Wg returns to a wild type-like pattern but often appeared distorted at the A/P boundary (Figure 8E), which we hypothesize could be due to misaligned compartments resulting from asymmetric growth (Figure 4). A similar elevation of Wg in the anterior and margin distortion occurs when *aspr-HA* is expressed only in the anterior (Figure S9D–H). These findings suggest that *aspr-HA* affects Wg during regeneration; however, Wg levels appear elevated rather than reduced, which is unexpected given the failure to reestablish multiple Wg target genes (*vg, Delta, ct, ac, al*) in the presence of Aspr-HA (Figures 5 and 6). Wg signaling depends on proper epithelial polarity; one model proposes that Wg is produced at the apical surface and subsequently trafficked to the basal side, where receptor engagement activates downstream signaling.^59,60^ Given that Aspr localizes to the apical membrane and within EVs situated above this surface (Figure 7 and Figure S7), we hypothesize that Aspr may modulate Wg availability by sequestering the ligand apically. Such sequestration could diminish the pool of Wg accessible for signaling and/or restrict its transcytosis to the basal surface.

**Figure 8.**
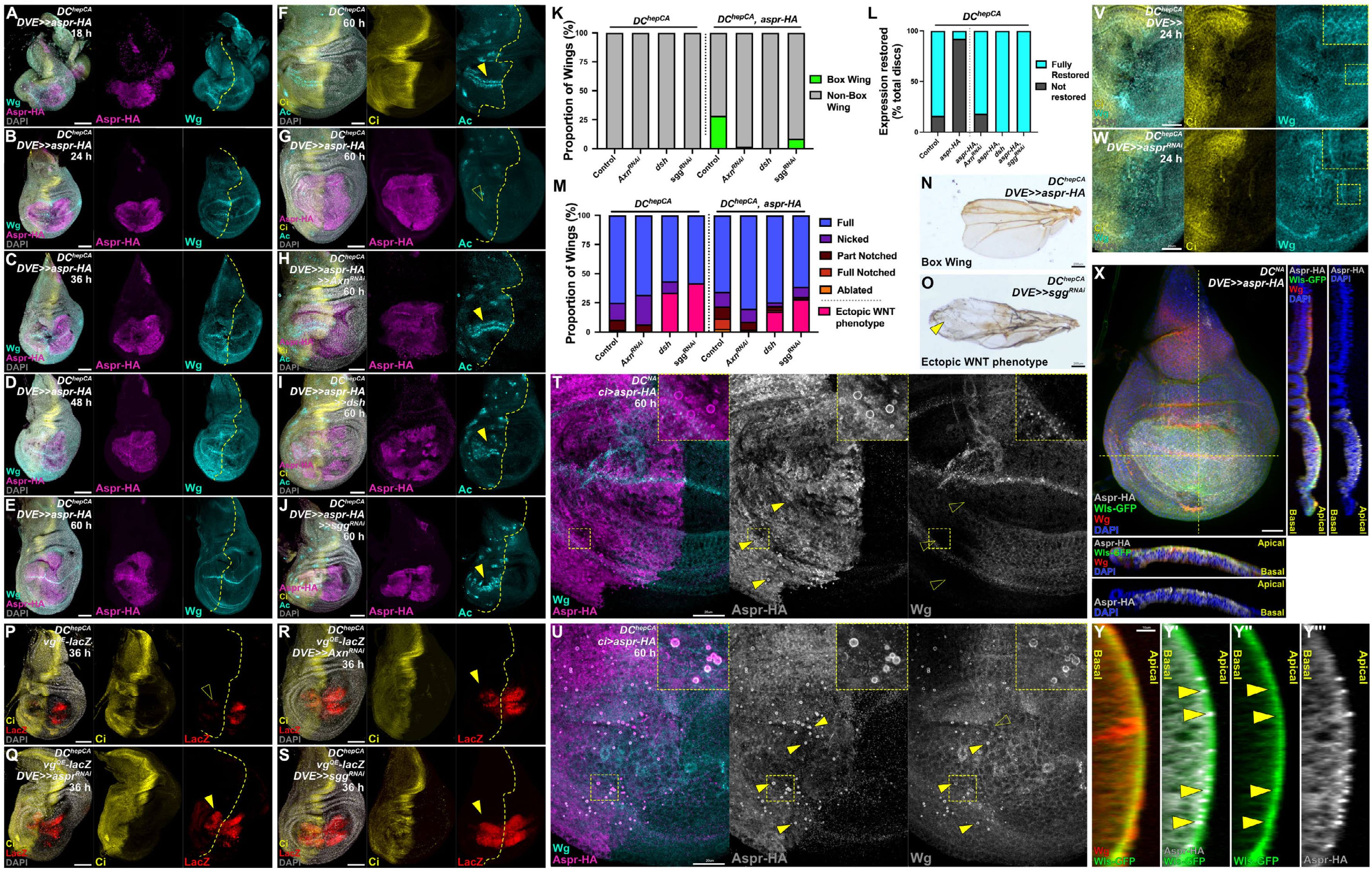
Aspr attenuates WNT signaling during regeneration. (**A–E**) Time course of Wg (cyan) in ablated discs (*DC^hepCA^, DVE>>GAL4*) expressing *aspr-HA* (magenta), 18–60 h AHS. Ci (yellow) marks anterior and the A/P boundary (dotted line). From 18–48 h, Wg signal appears increased anteriorly; by 60 h the anterior bias is lost but Wg patterning is distorted across the margin. (**F–J**) Ablated discs (*DC^hepCA^, DVE>>GAL4 )* at 60 h post-ablation, stained for Achaete (Ac, cyan) and Ci (yellow). Control discs (F) restore Ac expression (arrowhead), while *aspr-HA* blocks Ac restoration (G, open arrowhead). Co-expression of *aspr-HA* with WNT pathway activators *Axn^RNAi^* (H), *UAS-dsh* (I), or *sgg^RNAi^* (J) restores Ac (arrowheads). (**K**) Quantification of wing phenotypes from (F-J). Ectopic *aspr-HA* induces a box wing phenotype in 39% of wings. This is rescued by WNT activation: 1.7% with *Axn^RNAi^*, 0% with *dsh*, and 9.5% with *sgg^RNAi^*. Sample sizes: *DVE>>y^RNAi^*, n=275, *Axn^RNAi^*, n=255, *dsh*, n=122, *sgg^RNAi^*, n=146; *DVE>>aspr-HA* with *y^RNAi^*, n=243, *Axn^RNAi^*, n=119, *dsh*, n=63, *sgg^RNAi^*, n=288. (**L**) Quantification of Ac restoration in discs from (F-J). Ectopic Aspr-HA reduces Ac restoration to 8% of discs, whereas co-activation of Wg signaling restores Ac in 81% (*Axn^RNAi^*) and 100% (*dsh* and *sgg^RNAi^*). Sample sizes: GFP, n=56, *aspr-HA*, n=38, *aspr-HA*+*Axn^RNAi^*, n=11, *aspr-HA*+*dsh*, n=3, *aspr-HA*+*sgg^RNAi^*, n=2. (**M**) Quantification of regeneration outcomes from experiments in (K), categorized by severity of tissue loss. WNT activation alone produces a distinct “ectopic WNT” phenotype (pink), shown in (O). Co-expression with *aspr-HA* slightly reduces this phenotype. (**N–O**) Representative adult wing phenotypes from ablated discs (*DC^hepCA^, DVE>>GAL4)*. (N) Box wing phenotype from *aspr-HA*. (O) Ectopic WNT phenotype with mis-patterning and extra bristles. (**P–S**) Ablated discs (*DC^hepCA^, DVE>>GAL4)* at 36 h post-ablation bearing *vg^QE^-lacZ* and expressing RNAi for *aspr* (Q), *Axn* (R), or *sgg* (S). In control discs (P), *lacZ* is absent in the anterior (open arrowhead). All RNAi conditions restore anterior *lacZ* expression (arrowheads). (**T–U**) Wg (cyan) and Aspr-HA (magenta) colocalization at 60 h AHS. In unablated discs expressing *aspr-HA*, (*DC^NA^*, *ci-GAL4*) (T), Aspr-HA EVs (arrowheads) do not colocalize with Wg (open arrowheads). After damage (*DC^hepCA^, ci-GAL4)* (U), Wg is detected at Aspr-HA vesicles (arrowheads). Yellow dotted box indicates region shown at higher magnification insets. (**V–W**) Ablated discs (*DC^hepCA^, DVE>>GAL4)* at 24 h post-ablation stained for Wg (Cyan) and Ci (yellow), with knockdown of *aspr* (W), or control disc (V). Yellow dashed outline shows higher magnification inset. The loss of *aspr* reduces appearance of apical Wg puncta. (**X**) Unablated (*DC^NA^, DVE>>GAL4*) discs with the Wls-GFP reporter (green), expressing *aspr-HA* (grey) and stained for Wg (red). DAPI (blue) labels nuclei. Yellow dotted line indicates imaging planes shown in cross sections. Aspr-HA overlaps Wls staining within disc cells at the apical surface. (**Y-Y’’’**) High magnification transverse sections of disc in (X), showing the overlap of Wg with Wls (Y). and Wls with Aspr-HA (Y’-Y’’’). Yellow arrowheads indicate Aspr-associated EVs. All scale bars = 50 μm unless otherwise noted. See also Figures S8–S9. Full genotypes in Supplementary Genotypes.

If this model is correct, then activating Wg signaling downstream of the receptor should rescue the reestablishment of Wg target genes caused by sustained Aspr. To test this, we activated Wg signaling downstream of the receptor using three manipulations: knockdown of *Axin* (*Axn*) or *shaggy* (*sgg*), or overexpression of *disheveled* (*dsh*). We assessed expression of the Wg target *ac* as a readout. Normally, *ac* is restored in nearly all regenerating discs at 60 h AHS (Figure 8F) but is blocked by *aspr-HA* (Figure 8G). However, the activation of Wg signaling by any of the three manipulations fully rescues *ac* at 60 h AHS despite persistent *aspr-HA* expression (Figures 8H–J and 8L), confirming that Aspr can regulate Wg signaling at the level of the Wg ligand or receptor. Each manipulation alone also increases *ac* expression in both undamaged and regenerating discs (Figures S9I–N), with particularly strong effects when *dsh* is overexpressed (Figures S9J and M). *aspr-HA* mildly reduces these effects in undamaged discs (Figures S9O–R), suggesting that Aspr can temper ectopic Wg signaling to some degree even in the absence of injury. Importantly, these rescue experiments also significantly reduce the incidence of the “box wing” phenotype in regenerated wings (Figure 8K) and improves overall regenerative outcomes (Figure 8M). These manipulations, particularly the stronger *Dsh* overexpression and *sgg* knockdown, also result in a phenotype characterized by malformation and ectopic bristles within the wing blade (Figures 8M–O and S9S–T), suggesting these manipulations can go beyond rescuing the suppressive effects of *aspr-HA*, resulting in an “ectopic WNT” phenotype. This phenotype also occurs in undamaged discs at an equivalent frequency (Figure S9Y–AA) and therefore is unlikely to interfere with our interpretations in the context of regeneration.

These experiments are performed in the context of ectopic *aspr-HA* expression. To confirm that Wg signaling is regulated by Aspr endogenously, we knocked down *aspr* during regeneration and examined *vg*, a Wg responsive target gene^61^ that is present at the regenerative time point when Aspr is normally expressed, and that we know is affected by Aspr; indeed, we demonstrated that ectopic Aspr not only limits *vg* expression after damage (Figure 5H-H’’), but it can also blunt the growth induced by ectopic *vg* during regeneration (Figure 5M–O). Previous work has established that Vg and Wg cooperate in a growth-promoting feedback loop during development,^41,62–66^ and thus we hypothesize that if a similar dynamic occurs during regeneration, the addition of Aspr would interfere with this Vg-Wg-growth axis, resulting in the growth suppression we observe. To monitor *vg* we used *vg^QE^*, the vestigial quadrant enhancer that depends on Wg signaling for its activity,^38^ as a transcriptional readout. Under normal conditions, *vg^QE^* is not reactivated in the anterior pouch by 36 h AHS (Figure 8P). However, *aspr* knockdown leads to premature *vg^QE^* reactivation (Figure 8Q), supporting the model that Aspr delays Wg-dependent re-initiation of *vg*. Similarly, knockdown of *Axn* or *sgg* produces the same early-activation phenotype (Figures 8R–S), further strengthening this hypothesis. No alterations in *vg^QE^* activity were detected with these manipulations during development (Figures S9U–S9X).

Finally, we asked whether Aspr and Wg physically colocalize during regeneration. The anti-Aspr antibody was too weak to detect endogenous co-localization with Wg, and thus we examined Wg in aspr-HA expressing discs. In undamaged discs, even with optimized extracellular detection of Wg (see STAR Methods), Wg and Aspr showed no colocalization (Figure 8T). However, upon damage Wg now clearly associates with Aspr-positive EVs at the apical surface (Figure 8U). Notably, Wg localizes both on the surface and within these EVs (Figure S9AC), consistent with its presence as vesicular cargo. Supporting this interpretation, Wg staining is lost in non-permeabilized samples, indicating that its detectable epitope is internal rather than surface exposed (Figure S9AB). Moreover, *aspr* knockdown reduces the apical Wg-positive puncta normally observed in the tissue (Figure 8V–W), consistent with Aspr contributing to Wg trafficking or localization during regeneration. This idea is further supported by the colocalization of Aspr with Wntless (Wls-GFP), a well-established chaperone required for Wg secretion,^52^ at the apical surface of disc cells (Figure 8X–Y’’’). Because Aspr is enriched apically above the Crb marker (Figure 7G–H’’’) and shows minimal overlap with membrane-associated GFP within EVs (Figure S7Q–R’), we hypothesize that Wg resides within Aspr-associated EVs that exhibit several hallmarks of exosome-like vesicles, although their precise identity remains to be determined. Together, these observations support a model in which Aspr modulates Wnt signaling during tissue repair by sequestering Wg within these EVs.

Thus, we hypothesize that Aspr regulates the timing of Wg signaling targets by modulating Wg ligand availability (Figure 9). Upon damage, *aspr* is initially expressed via its DRMS enhancer in blastema cells alongside *wg* and other secreted factors. During these early regeneration stages, Aspr may facilitate Wg trafficking to promote balanced compartment growth via *vg*, while restraining activation of late patterning genes such as *bs* and *ac*. After 24 h, Aspr levels decline, allowing Wg signaling to reinitiate differentiation programs to promote proper repatterning. Artificially maintaining Aspr beyond this window sequesters Wg apically in EVs, disrupting growth balance and repatterning, resulting in poor regeneration and the characteristic “box wing” phenotype. Conversely, *aspr* knockdown causes premature target activation, further disrupting the precise regenerative sequence required for full wing restoration.

**Figure 9.**
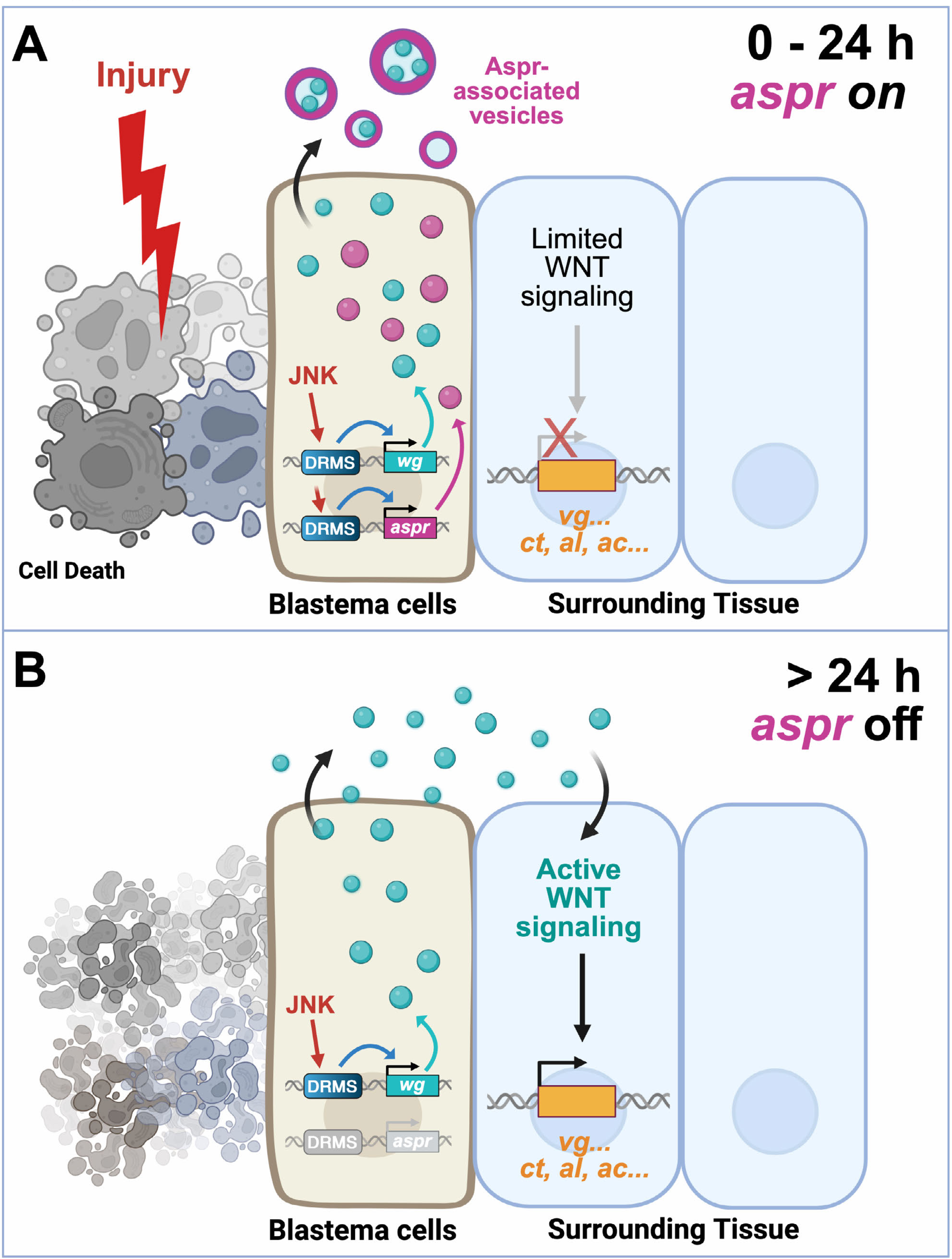
Functional model: Aspr temporally regulates WNT signaling during regeneration. (**A–B**) Schematic model proposing that (A) early after injury (first ∼24 h), dying cells induce blastema formation in adjacent tissue via JNK activity, activating DRMS enhancers to drive *aspr* and *wg* expression. Aspr localizes to vesicles that sequester Wg, limiting its access to neighboring cells. (B) After ∼24 h, Aspr levels fall, permitting Wg to engage receptors and activate WNT signaling, which triggers cell-fate gene expression (e.g., *vg, ac, ct, al)* and re-patterning only after sufficient growth has occurred, thereby coordinating growth and patterning during regeneration.

## Discussion

### Aspr regulates regenerative WNT signaling

Regeneration reactivates developmental programs, yet it often must occur within mature tissues, which poses unique challenges. Temporal coordination is particularly important, as fate specification must follow tissue regrowth to ensure proper repatterning.^1,67^ Our findings identify *aspr* as a primarily regeneration-specific gene that accomplishes this by attenuating Wg signaling during early regeneration. *aspr* appears dispensable during development but is activated by damage-induced JNK signaling via a DRMS enhancer,^68^ where it functions to properly time WNT-dependent patterning genes in regenerating tissue, potentially via sequestration of the Wg ligand in apically secreted extracellular EVs. This mechanism differs from previously described temporal regulators of regeneration, such as Ets21C and Brat, which act at the level of transcription and translation.^31,69^ By contrast, Aspr potentially modulates ligand availability, demonstrating an alternative strategy by which the sequential steps of a regeneration program are maintained. We find that artificially maintaining Aspr disrupts this timing, delaying the restoration of WNT target genes, and leading to defective patterning and disproportional/restricted growth.

These effects are rescued by pathway activation downstream of the receptor, supporting a model in which Aspr functions at or above the level of WNT ligand/receptor interaction to negatively regulate Wg signaling during early regeneration. It therefore joins factors such as Evi/Wntless,^52^ Porcupine^70,71^, SWIM^72^, and the heparan sulfate proteoglycans Dally and Dally-like^73^, which regulate Wg signaling by controlling overall ligand availability during disc development.^57^ Thus, exploring how these factors function in the context of regeneration represents an important direction for future research. Aspr is unlikely to completely block Wg signaling, but rather Aspr may fine-tune Wg signaling following injury; its ability to antagonize WNT activity when co-expressed with activated pathway components, even in undamaged discs (Figure S9P-S9R), suggests it may buffer excess signaling, which is particularly important in regenerative contexts in which signaling is inherently more variable. In regenerating wing discs, Wg is strongly induced and broadly expressed at early stages before its expression narrows and declines^69^, while in a developmental context different levels of Wg/Wnt activity are known to elicit distinct cellular responses ranging from survival to proliferation and differentiation.^74^ These features are consistent with a model in which Aspr functions as a regeneration specific modulator that keeps Wg signaling within an optimal range, supporting initial proportional growth while preventing premature differentiation.

Interestingly, the effects of maintaining *aspr* expression are more impactful in the anterior compartment. Ectopic Aspr preferentially disrupts anterior patterning, growth, and cell fate markers, and anterior proliferation is more sensitive to its expression. While Aspr is broadly induced in the pouch during wounding, this compartment-specific effect may reflect intrinsic differences in signaling dynamics, baseline proliferation rates, or the influence of specific regulators that are unique to each compartment. The observation that Aspr protein is not equally detected in the anterior versus posterior compartment of the wing disc when ectopically expressed under pouch-wide UAS control, despite transcripts being uniformly distributed, also raises the possibility that compartment-specific post-transcriptional mechanisms influence Aspr. Such asymmetries could reflect differential protein stability, localized degradation pathways, or compartment-specific modulation of secretion, trafficking, or extracellular retention. Alternatively, Aspr may require cofactors or interacting partners that are present at different levels across compartments, resulting in localized stabilization, enhanced turnover or differences in functionality. Thus, the wing disc’s intrinsic anterior–posterior heterogeneity may shape the distribution and activity of Aspr through several potential mechanisms, emphasizing the importance of understanding how tissue architecture and compartmental identity influence regenerative capacity. Although compartment-specific identity genes are thought to be largely maintained or quickly reestablished during regeneration,^69,75–77^ additional studies such as those utilizing single cell sequencing,^31^ may help to elucidate how compartment identity influences regenerative mechanisms and outcomes.

### EV-based control over regenerative timing

During development of the wing disc, Wg is secreted apically and trafficked to the basolateral membrane for receptor interaction.^26,57,59,78–80^ Our results suggest that in regeneration, Aspr delays this process by associating with Wg in vesicles that remain apical and separate from the signaling-competent basolateral domains. This may block WNT signaling by physically preventing ligand-receptor interaction, consistent with findings that altering Wg ligand internalization can block signaling.^57,81–83^ Once *aspr* expression declines after 24 hours, Wg becomes available for signaling, initiating cell fate specification at the appropriate time. Alternatively, Aspr may alter the secretion or modification of Wg to alter its ability to signal in other ways, since alternative trafficking is known to alter secretion rates, location and post-translational modifications of ligands.^57^ Nevertheless, this EV-based mechanism raises several questions. Is Wg encapsulation actively regulated, or a passive consequence of altered trafficking? What is the fate of Aspr vesicles? Do they carry other morphogens, such as Dpp, or regulatory factors involved in regeneration? The suppression of Dpp target *al* by ectopic Aspr, and the known crosstalk between Wg and Dpp,^84–86^ suggests broader regulatory functions that remain to be explored. The localization of Wg to Aspr-containing vesicles during regeneration and not development suggests that additional injury-induced factors may mediate cargo trafficking into these EVs. Given that both Wg and Aspr are regulated by DRMS enhancers,^22^ and are co-expressed in “secretory zone” cells of the blastema,^31^ it is possible that other factors involved in EV formation or cargo selection are similarly regulated. Therefore, systematic identification of DRMS enhancer targets may uncover additional components or cargoes of EVs that help coordinate regeneration through this mechanism.

### Broader implications and future perspectives

This work identifies *aspr* as a novel regeneration-specific gene that regulates tissue patterning by modulating WNT ligand availability via vesicle-based sequestration. These findings underscore the importance of temporal control mechanisms in regeneration and highlight vesicle trafficking as an underexplored mode of signal regulation.

Beyond *Drosophila*, EV-mediated intercellular communication is increasingly recognized as critical in cancer and immune biology.^87–90^ Our findings suggest that analogous mechanisms may exist in injury and regeneration contexts, and could be harnessed to enhance tissue repair. Future work should focus on the molecular determinants of Wg loading into Aspr vesicles, the identity of additional vesicle cargo, and whether other DRMS-induced genes in blastema cells are regulated via similar mechanisms. Exploring whether vertebrate functional homologs of *aspr* exist, and whether they influence WNT signaling during injury, may open new avenues for regenerative therapies. Ultimately, understanding how tissues control not just which genes are expressed, but also when they are activated, may be essential to unlock the full potential of regenerative biology.

### Limitations of the Study

In this study, we identify a regeneration-specific role for Aspr, a largely uncharacterized factor induced by damage in the wing disc. Our findings suggest that Aspr may act through the formation or modulation of extracellular vesicles (EVs) that influence Wg ligand availability; however, the precise nature, origin, and functional contribution of these EVs remain incompletely understood. Additional work using high-resolution imaging and biochemical EV isolation will be required to determine how these structures affect Wg secretion or trafficking, and to characterize their endogenous formation and behavior without relying on Aspr overexpression.

Furthermore, it is not yet known whether these EV-like structures contain additional molecular cargo or impact signaling pathways beyond WNT. Finally, our ability to study endogenous Aspr is limited by the low sensitivity of the antibody generated for this work. Development of an endogenously tagged Aspr allele will be critical for defining its localization, dynamics, and function during regeneration.

## Supporting information

STAR Methods

## Resource Availability

### Lead contact

Further information and requests for resources and reagents should be directed to and will be fulfilled by the lead contact, Robin Harris

### Materials availability

Fly lines generated or used in this work are available upon request; further details regarding stock genotypes are also provided in the Supplementary Genotypes file. For more detailed protocols, please contact the first author, and any further information required to reproduce the experiments can be obtained by contacting the corresponding author.

### Data Availability

All primary data supporting these findings are included within the manuscript or in the supplemental materials.

## Acknowledgements

The authors would like to thank Dr. Iswar Hariharan and Dr. David Bilder of UC Berkeley, Dr. Mirka Uhlirova of the University of Cologne, Dr. Seth Bliar of the University of Wisconsin, Dr. Kirsten Guss of Dickinson College, Dr. Baotong Xie of Oregon Health & Science University, Dr. Brian Gebelein of Cincinnati’s Children’s, Dr. Sonsoles Campuzano of the Centro de Biología Molecular Severo Ochoa, Dr. Gerard Campbell of the University of Pittsburgh for stocks and reagents. Current and former members of the Harris lab for feedback, and the Bloomington Stock Center, Molecular Instruments, and Developmental Studies Hybridoma Bank for stocks and reagents. This research has received no external funding.

## Author Contributions

Si Cave, Conceptualization, Methodology, Investigation, Data Curation, Formal Analysis, Validation, Writing – Original Draft, Writing – Review and Editing, Visualization; Maksym Dankovsky, Investigation, Data Curation, Formal Analysis, Validation, Writing – Review and Editing, Visualization; Jordan Hieronymus, Investigation, Data Curation, Formal Analysis, Validation, Writing – Review and Editing, Visualization; Manashi Sonowal, Conceptualization, Investigation Data Curation, Formal Analysis, Validation, Writing – Original Draft, Chloe Van Hazel, Investigation Data Curation, Validation, Resources; Petra Fromme, Supervision, Conceptualization, Resources, Robin Harris, Supervision, Conceptualization, Methodology, Investigation, Data Curation, Formal Analysis, Funding Acquisition, Writing – Original Draft, Writing – Review and Editing, Visualization, Project Administration.

## Supplemental Information

Document S1. Figures S1–S9, Supplemental Genotypes document.

