## Supplementary material for "Asperous coordinates regenerative timing by regulating damage-induced WNT Signaling": STAR Methods

**Key Resources Table**

| REAGENT or RESOURCE | SOURCE | IDENTIFIER |
| --- | --- | --- |
| Antibodies | | |
| Rat anti-Cubitus interruptus (Ci) | DSHB | Cat# [2A1](https://dshb.biology.uiowa.edu/2A1);  RRID: [AB_2109711](https://www.antibodyregistry.org/AB_2109711) |
| Mouse anti- Matrix Metalloproteinase 1 (Mmp1) C-terminus | DSHB | Cat# [14A3D2](https://dshb.biology.uiowa.edu/14A3D2);  RRID: [AB_579782](https://www.antibodyregistry.org/AB_579782) |
| Mouse anti- Matrix Metalloproteinase 1 (Mmp1) catalytic domain | DSHB | Cat# [3A6B4](https://3A6B4); RRID:[AB_579780](https://www.antibodyregistry.org/AB_579780) |
| Mouse anti-Wingless (Wg) | DSHB | Cat# [4D4](https://dshb.biology.uiowa.edu/4D4); RRID:[AB_528512](https://www.antibodyregistry.org/AB_528512) |
| Mouse anti-Nubbin (Nub) | DSHB | Cat# [nub2D4](https://dshb.biology.uiowa.edu/Nub-2D4); RRID:[AB_2722119](https://www.antibodyregistry.org/AB_2722119) |
| Mouse anti-LacZ | DSHB | Cat# [40-1a](https://dshb.biology.uiowa.edu/40-1a); RRID:[AB_528100](https://www.antibodyregistry.org/AB_528100) |
| Mouse anti-Achaete (Ac) | DSHB | Cat# [anti-achaete](https://dshb.biology.uiowa.edu/anti-achaete) RRID:[AB_528066](https://www.antibodyregistry.org/AB_528066) |
| Mouse anti-Cut (Ct) | DSHB | Cat# [2B10](https://dshb.biology.uiowa.edu/2B10); RRID:AB_[528186](https://www.antibodyregistry.org/AB_528186) |
| Mouse anti-Delta (Dl) extracellular | DSHB | Cat# [C594.9B](https://dshb.biology.uiowa.edu/C594-9B); RRID:[AB_528194](https://www.antibodyregistry.org/AB_528194) |
| Mouse anti-Discs large (Dlg) | DSHB | Cat# [4F3](https://dshb.biology.uiowa.edu/4F3-anti-discs-large); RRID:[AB_528203](https://www.antibodyregistry.org/AB_528203) |
| Rabbit anti-Hemagglutinin (HA) | Cell Signaling Technology | Cat# [3724](https://www.cellsignal.com/products/primary-antibodies/ha-tag-c29f4-rabbit-mab/3724?srsltid=AfmBOor-QrjNPPo6pTnPaJiRC2DTDF4TSGBYApLrTFJIKadxH4KxJbah); RRID:[AB_1549585](https://www.antibodyregistry.org/AB_1549585) |
| Mouse anti-Hemagglutinin (HA) | Cell Signaling Technology | Cat# [2367](https://www.cellsignal.com/products/primary-antibodies/ha-tag-6e2-mouse-mab/2367); RRID:[AB_10691311](https://www.antibodyregistry.org/AB_10691311) |
| Rabbit anti-cleaved Death caspase-1 (Dcp-1) (Asp215) | Cell Signaling Technology | Cat# [9578](https://www.cellsignal.com/products/primary-antibodies/cleaved-drosophila-dcp-1-asp215-antibody/9578); RRID:[AB_2721060](https://www.antibodyregistry.org/AB_2721060) |
| Mouse anti-P-histone H3 (Ser10) | Cell Signaling Technology | Cat# [9706](https://www.cellsignal.com/products/primary-antibodies/phospho-histone-h3-ser10-6g3-mouse-mab/9706); RRID:[AB_331748](https://www.antibodyregistry.org/AB_331748) |
| Rabbit anti-Phospho-SMAD1/5 (Ser463/465) | Cell Signaling Technology | Cat# [9516](https://www.cellsignal.com/products/primary-antibodies/phospho-smad1-5-ser463-465-41d10-rabbit-mab/9516); RRID:[AB_491015](https://www.antibodyregistry.org/AB_491015) |
| Rat anti-DE-Cadherin | DSHB | Cat# DCAD2;  RRID: [AB_528120](https://www.antibodyregistry.org/AB_528120) |
| Chicken anti-Green Fluorescent Protein (GFP) | Abcam | Cat# [ab13970](https://www.abcam.com/en-us/products/primary-antibodies/gfp-antibody-ab13970?srsltid=AfmBOoriysu2rBAcwBo9Lm8IMB-u9cL8JZ1QDn-1Xl90wT9s9223oKEu); RRID:[AB_300798](https://www.antibodyregistry.org/AB_300798) |
| Mouse anti-beta Actin (β-Act) | Abcam | Cat# [ab8442](https://www.abcam.com/en-us/products/primary-antibodies/beta-actin-antibody-mabcam-8224-loading-control-ab8224?srsltid=AfmBOoqV3X9QIP8rXevWdqadd3G558nN2nVdICOtdENZTFy39081Gg1a#drawerView=quickview);  RRID:[AB_449644](https://www.antibodyregistry.org/AB_449644) |
| Rabbit anti-Asperous (Aspr) | Pacific Immunology | Custom; N/A |
| Rabbit anti-Vestigial (Vg) | Dr. Kirsten Guss (Gift), *Williams et al.^83?^* | N/A |
| Mouse anti-Blistered (Bs) | Dr. Seth Blair (Gift),  *Affolter et al.^84^* | N/A |
| Rabbit anti-Salm | Dr. Baotong Xie and Dr. Brian Gebelein (Gift), *Xie et al.^85^* | N/A |
| Rat anti-Aristaless (Al) | Dr. Gerard Campbell (Gift), *Campbell et al.^86^* | N/A |
| Goat anti-Rat Alexa 647 | Invitrogen | Cat # [A21247](https://www.thermofisher.com/antibody/product/Goat-anti-Rat-IgG-H-L-Cross-Adsorbed-Secondary-Antibody-Polyclonal/A-21247)  RRID: [AB_141778](https://www.antibodyregistry.org/AB_141778) |
| Goat anti-Rat Alexa 555 | Invitrogen | Cat# [A21434](https://www.thermofisher.com/antibody/product/Goat-anti-Rat-IgG-H-L-Cross-Adsorbed-Secondary-Antibody-Polyclonal/A-21434);  RRID: [AB_2535855](https://www.antibodyregistry.org/AB_2535855) |
| Goat anti-Rat Alexa 488 | Invitrogen | Cat# [A11006](https://www.thermofisher.com/antibody/product/Goat-anti-Rat-IgG-H-L-Cross-Adsorbed-Secondary-Antibody-Polyclonal/A-11006);  RRID: [AB_2534074](https://www.antibodyregistry.org/AB_2534074) |
| Goat anti-Mouse Alexa 647 | Invitrogen | Cat# [A21236](https://www.thermofisher.com/antibody/product/Goat-anti-Mouse-IgG-H-L-Highly-Cross-Adsorbed-Secondary-Antibody-Polyclonal/A-21236);  RRID: [AB_2535805](https://www.antibodyregistry.org/AB_2535805) |
| Goat anti-Mouse Alexa 555 | Invitrogen | Cat# [A32727](https://www.thermofisher.com/antibody/product/Goat-anti-Mouse-IgG-H-L-Highly-Cross-Adsorbed-Secondary-Antibody-Polyclonal/A32727);  RRID: [AB_2633276](https://www.antibodyregistry.org/AB_2633276) |
| Goat anti-Mouse Alexa 488 | Invitrogen | Cat# [A32723](https://www.thermofisher.com/antibody/product/Goat-anti-Mouse-IgG-H-L-Highly-Cross-Adsorbed-Secondary-Antibody-Polyclonal/A32723);  RRID: [AB_2633275](https://www.antibodyregistry.org/AB_2633275) |
| Goat anti-Rabbit Alexa 647 | Invitrogen | Cat# [A21245](https://www.thermofisher.com/antibody/product/Goat-anti-Rabbit-IgG-H-L-Highly-Cross-Adsorbed-Secondary-Antibody-Polyclonal/A-21245); RRID: [AB_2535813](https://www.antibodyregistry.org/AB_2535813) |
| Goat anti-Rabbit Alexa 555 | Invitrogen | Cat# [A32732](https://www.thermofisher.com/antibody/product/Goat-anti-Rabbit-IgG-H-L-Highly-Cross-Adsorbed-Secondary-Antibody-Polyclonal/A32732);  RRID: [AB_2633281](https://www.antibodyregistry.org/AB_2633281) |
| Goat anti-Rabbit Alexa 488 | Invitrogen | Cat# [A11006](https://www.thermofisher.com/antibody/product/Goat-anti-Rat-IgG-H-L-Cross-Adsorbed-Secondary-Antibody-Polyclonal/A-11006);  RRID: [AB_2534074](https://www.antibodyregistry.org/AB_2534074) |
| Goat anti-Chicken Alexa 488 | Invitrogen | Cat# [A11039](https://www.thermofisher.com/antibody/product/Goat-anti-Chicken-IgY-H-L-Secondary-Antibody-Polyclonal/A-11039);  RRID: [AB_2534096](https://www.antibodyregistry.org/AB_2534096) |
| Goat anti-Guinea Pig Alexa 647 | Invitrogen | Cat# [A21450](https://www.thermofisher.com/antibody/product/Goat-anti-Guinea-Pig-IgG-H-L-Highly-Cross-Adsorbed-Secondary-Antibody-Polyclonal/A-21450); RRID: [AB_2535867](https://www.antibodyregistry.org/AB_2535867) |
| Bacterial and virus strains | | |
| XL10-Gold Ultracompetent Cells | Agilent | Cat# [200314](https://www.agilent.com/en/product/mutagenesis-cloning/competent-cells-competent-cell-supplies/competent-cells-for-difficult-cloning/xl10-gold-ultracompetent-cells-233087) |
| Chemicals, peptides, and recombinant proteins | | |
| ProLong Diamond Antifade Mountant | Life Technologies | Cat# [P36965](https://www.thermofisher.com/order/catalog/product/P36965) |
| Hybridization Chain Reaction (HCR^TM^) RNA-FISH probe sets, amplifiers, and buffers | Molecular Instruments | HCR RNA-FISH Bundles |
| DAPI | ThermoFisher | Cat# [D1306](https://www.thermofisher.com/order/catalog/product/D1306) |
| Paraformaldehyde (PFA) | ThermoFisher | Cat# [28909](https://www.thermofisher.com/order/catalog/product/28906) |
| Fisher Chemical^TM^ Permount^TM^ Mounting Medium | Fisher Scientific | Cat# [SP15-500](https://www.fishersci.com/shop/products/fisher-chemical-permount-mounting-medium-2/SP15100?ef_id=Cj0KCQjwxo_CBhDbARIsADWpDH7bxZmp4uj9tbHJzqavjS8JvUdbd4jdjNKMf7MaeaGENJgGhwCG9woaAqwtEALw_wcB:G:s&ppc_id=PLA_goog_2086145683_86816103982_SP15100__411323788695_7402561504943389543&ev_chn=shop&s_kwcid=AL!4428!3!411323788695!!!g!859093046079!&gad_source=1&gad_campaignid=2086145683&gbraid=0AAAAADu8mFy7NRts9XyfWLf_tJcWlVg3l&gclid=Cj0KCQjwxo_CBhDbARIsADWpDH7bxZmp4uj9tbHJzqavjS8JvUdbd4jdjNKMf7MaeaGENJgGhwCG9woaAqwtEALw_wcB) |
| Triton X-100 | Sigma-Aldrich | Cat# [T8787](https://www.sigmaaldrich.com/US/en/product/sigma/t8787?srsltid=AfmBOoqDypFIFWMRTsFcnxA1xtGrYrbqxUmJMq_6s_2T8mMiS3Zz8RrY) |
| Tween-20 | Sigma-Aldrich | Cat# [P1379](https://www.sigmaaldrich.com/US/en/product/sial/p1379?srsltid=AfmBOooHtMGDpDENldVXVrz5yhOur27aDYKJg6A1RK0DW-uOw4vqEpGL) |
| Tris-EDTA (TE) Buffer | Thermo Fisher Scientific | Cat# [BP2473500](https://www.fishersci.com/shop/products/tris-edta-1x-solution-ph-8-0-molecular-biology-fisher-bioreagents-3/BP2473500#?keyword=fisher) |
| 20X Sodium Chloride Sodium Citrate (SSC) | G-Biosciences | Cat# [R019](https://www.gbiosciences.com/Buffers-Reagents-Chemicals/Molecular-Biology-Related-Buffers-Chemicals/SSC-Buffer-20X) |
| Non-Goat Serum (NGS) | Abcam | Cat# [AB7481](https://www.abcam.com/en-us/products/blood-derived-fluids/normal-goat-serum-ab7481?srsltid=AfmBOorPuyJRxm5939dlRa2KVGcL-c8jDRcb0K5C5dYeyIjI48YaT45C) |
| Insect PopCulture | Sigma Aldrich^TM^ | Cat# [I7267](https://www.sigmaaldrich.com/US/en/product/sigma/i7267?srsltid=AfmBOoqOT1iIkwC3kvjGb2ONDYBYlEoDSwhfCDOPwbKlDhkq6gx6bGqQ) |
| Immun-Star Chemiluminescence Kit | Bio-Rad | Cat# [170-5010](https://www.bio-rad.com/en-us/sku/1705010-immun-star-goat-anti-mouse-gam-ap-detection-kit?ID=1705010) |
| PVDF Membrane | Bio-Rad/Millipore | Cat# [1620177](https://www.bio-rad.com/en-us/product/immun-blot-pvdf-membrane?ID=9f95f7dc-8ad2-4923-8eb4-be822b64e2f4) |
| Critical commercial assays | | |
| HCR™ RNA-FISH (v3.0) Kit | Molecular Instruments | [v3.0](https://store.molecularinstruments.com/new-kit/v3/rnafish) |
| Click-iT^TM^ Edu Cell Proliferation Kit for Imaging, Alexa Fluro^TM^ 555 Dye | Invitrogen | Cat# [10338](https://www.thermofisher.com/order/catalog/product/C10338) |
| Experimental models: Organisms/strains | | |
| *hs-flp; hs-p65; salm-LexADBD/TMBB dve>>Gal4 (DC^NA^ ^DVE^)* | Bloomington Drosophila Stock Center | BDSC:605701 |
| *hs-flp; hs-p65, lexAop-hepCA; salm-LexADBD/TM6B (DCh^epCA^ ^No Gal4^)* | Bloomington Drosophila Stock Center | BDSC:605694 |
| *hs-flp; hs-p65, lexAop-hepCA/Cyo; salm-LexADBD, dve>>Gal4/TM6C (DC^hepCA^ ^DVE^)* | Bloomington Drosophila Stock Center | BDSC:605695 |
| *patched-Gal4* | Bloomington Drosophila Stock Center | BDSC:44612 |
| *scalloped-Gal4* | Bloomington Drosophila Stock Center | BDSC:40674 |
| *pannier-Gal4* | Bloomington Drosophila Stock Center | BDSC:3039 |
| *hedgehog-Gal4* | Bloomington Drosophila Stock Center | BDSC:600186 |
| *UAS-RFP* | Bloomington Drosophila Stock Center | BDSC:30556 |
| *UAS-y^RNAi^* | Bloomington Drosophila Stock Center | BDSC:28710 |
| *UAS-hep^CA^* | Bloomington Drosophila Stock Center | BDSC:6406 |
| *UAS-JNK^DN^* | Bloomington Drosophila Stock Center | BDSC:6409 |
| *PCNA-GFP* | Bloomington Drosophila Stock Center | BDSC:25749 |
| *AP1-RFP* | Bloomington Drosophila Stock Center | BDSC:59011 |
| *Dpp-LacZ* | Bloomington Drosophila Stock Center | BDSC:8412 |
| *hep^-^* | Bloomington Drosophila Stock Center | BDSC:6761 |
| *Aspr^Mi(MIC)^* | Bloomington Drosophila Stock Center | BDSC:35864 |
| *UAS-Dsh-Myc* | Bloomington Drosophila Stock Center | BDSC:9453 |
| *Sgg^RNAi^* | Bloomington Drosophila Stock Center | BDSC:35364 |
| *Axn^RNAi^* | Bloomington Drosophila Stock Center | BDSC:31705 |
| *Crb-GFP* | Bloomington Drosophila Stock Center | BDSC:99495 |
| *Ct-Gal4* | Bloomington Drosophila Stock Center | BDSC:27327 |
| *UAS-vg* | Bloomington Drosophila Stock Center | BDSC:37296 |
| *Wls-GFP* | Bloomington Drosophila Stock Center | BDSC: 94900 |
| *UAS-CD8::GFP* | Bloomington Drosophila Stock Center | BDSC: 5130 |
| *Lgl^RNAi^* | Vienna Drosophila Resource Center | VDRC:51247 |
| *hs-p65/Cyo; salm-LexADBD/TMB6 (DC^NA No Gal4^)* | *Harris et al.^22^* | N/A |
| *hs-flp hs-p65, lexAop-gluR1Lc8; salm-LexADBD (DC^gluR1^ ^No Gal4^)* | *Klemm et al.23* | N/A |
| *UAS Aspr^RNAi M2^* | Dr. David Bilder (Gift),  Unpublished | N/A |
| *UAS-Aspr-FLAG.3xHA* | Drosophila Genomic Resource Center | Barcode: 1063862 |
| *UAS-GFPnls* | Dr. Iswar Hariharan (Gift) |  |
| *ci-Gal4* | Dr. Iswar Hariharan (Gift) | N/A |
| *UAS-egr* | Dr. Iswar Hariharan (Gift) | N/A |
| *Vg^QE^-LacZ* | Dr. Tin Tin Su (Gift),  *Kim et al.^87^* | N/A |
| *UAS-Aspr* | This paper | N/A |
| *UAS-Aspr-HA* | This paper | N/A |
| Oligonucleotides | | |
| Aspr cDNA Insertion | Berkely Drosophila Genome Project | LP13770 |
| Aspr Forward Primer *AGGGAATTGGGAATTCATGCAGCAGCTCCTGGTCG* | IDT DNA | N/A |
| Aspr Reverse Primer with HA | IDT DNA | N/A |
| White + Forward Primer  GCATCTCAAAAAAATGGTGGGCATAAT | IDT DNA | N/A |
| SV40 Reverse Primer  AGGTTCCTTCACAAAGATCC | IDT DNA | N/A |
| Recombinant DNA | | |
| pUAS-attB-Aspr-HA | Dr. Iswar Hariharan (Gift) | N/A |
| *aspr* HCR Probe | Molecule Instruments | Custom [NM_134528](https://www.ncbi.nlm.nih.gov/gene?Db=gene&Cmd=DetailsSearch&Term=33013) |
| *eGFP* HCR Probe | Molecule Instruments | Custom KC896842 |
| Software and algorithms | | |
| BioRender | BioRender | [https://biorender.com](https://www.biorender.com/) (academic license) |
| GraphPad Prism for Mac (Version 10.5.0) | GraphPad Software Inc. | <https://www.graphpad.com/features> |
| Affinity Photo | Serif | [https://affinity.serif.com/photo](https://affinity.serif.com/en-us/photo/) |
| Affinity Desigern | Seif | <https://affinity.serif.com/en-us/designer/> |
| Zeiss ZEN software | Zeiss | [https://www.zeiss.com](https://www.zeiss.com/corporate/en/home.html) |
| Fiji/ImageJ | NIH | <https://imagej.net/ij/> |
| AlphaFold v3 | DeepMind | [https://alphafold.ebi.ac.uk](https://alphafold.ebi.ac.uk/) |
| SignalP | DTU Bioinformatics | [https://services.healthtech.dtu.dk/service.php?SignalP](https://services.healthtech.dtu.dk/services/SignalP-6.0/) |
| Phobius | Stockholm University | [https://phobius.sbc.su.se](https://phobius.sbc.su.se/) |
| TargetP | DTU Bioinformatics | [https://services.healthtech.dtu.dk/service.php?TargetP](https://services.healthtech.dtu.dk/services/TargetP-2.0/) |
| Protter | ETH Zurich | [https://wlab.ethz.ch/protter/](https://wlab.ethz.ch/protter/start/) |
| DeepTMHMM | DTU Bioinformatics | <https://dtu.biolib.com/DeepTMHMM> |
| DeepLoc | DTU Bioinformatics | [https://services.healthtech.dtu.dk/service.php?DeepLoc](https://services.healthtech.dtu.dk/services/DeepLoc-2.1/) |
| UCSF ChimeraX1.9 | UCSF | <https://www.cgl.ucsf.edu/chimerax/> |
| PyMOL | Schrödinger | [https://pymol.org](https://pymol.org/) |
| Other | | |
| Mini-Strainer 0.40µM Filter | PluriSelect | Cat# [43-10040-40](https://www.pluriselect-usa.com/us/pluristrainer-mini-40-m-cell-strainer.html#size=27) |
| Zeiss AxioImager M2 with Apotome | Zeiss | N/A |
|  | GE Healthcare | N/A |
| Mini-PROTEAN Tetra Cell | Bio-Rad | Cat# [1658000](https://www.bio-rad.com/en-us/sku/1658000-mini-protean-tetra-cell-for-0-75-mm-gels?ID=1658000) |
| PowerPac Basic Power Supply | Bio-Rad | Cat# [1645050](https://www.bio-rad.com/en-us/sku/1645050-powerpac-basic-power-supply?ID=1645050) |
| eBlot Protein Trasnfer System | GenScript | Cat# [L00686C](https://www.genscript.com/eBlot-L1-protein-transfer-system.html) |
| Transgenic Injection Service | BestGene, Inc | N/A |

**Resource Availability**

**Lead Contact**

**Materials Availability**

Plasmids and transgenic *Drosophila* lines generated in this study, including the *UAS-Aspr* and *UAS-Aspr-HA* constructs, are available upon request from the lead contact. Newly generated antibodies are also available upon request.

**Data and Code Availability**

- All data reported in this study will be provided upon reasonable request from the lead contact
- This study did not generate the original code.
- Any additional information required to reanalyze the data reported in this paper is available upon request from the led contact.

**Experimental Model and Study Participant Details**

**Drosophila Stocks**

Stocks were maintained on standard dextrose-based medium containing 9.3 g agar, 32 g yeast, 61 g cornmeal, 129 g dextrose, and 14 g tegosept per liter of distilled water. Flies were raised at temperatures ranging from 18°C to 30°C, depending on the experimental requirements, under a 12-hour light/dark cycle. Genotype details for each figure panel are provided in the Supplementary Genotypes file.

**Method Details**

**Genetic Ablation using the DUAL Control System**

Ablations were induced using the DUAL Control (Duration and Location Control, DC) genetic system as previously described (Harris et al., 2020). Larvae were heat-shocked at 37°C for 45 minutes, followed by recovery periods at 25°C ranging from 12 to 60 hours, depending on the experiment.

**Immunohistochemistry and Microscopy**

Larvae were inverted in PBS, fixed with 4% paraformaldehyde (PFA) for 20 minutes, washed 3 times with 0.1% PBST (1x PBS + 0.1% Triton-X), permeabilized in 0.3% PBST. Samples were then blocked with 0.1% PBST with 5% normal goat serum (NGS) for 30 minutes. Tissues were incubated overnight at 4C. The following day, fluorescent secondary antibodies (1:500) were applied for 4 hours at room temperature. Inverted carcasses were dissected and mounted with ProLongTM Diamond Antifade Mountant from Life Technologies Company. Images were taken on a Zeiss AxioImager M2 with Apotome. All images were taken at 20X, unless otherwise indicated. Primary antibodies obtained from Developmental Studies Hybridoma Bank (DSHB) include: rat anti-Cubitius interruptus (Ci) (2A1, 1:10), mouse anti-Matrix metalloproteinase 1 (Mmp1) C-terminus (14A3D2, 1:500), mouse anti-Mmp1 catalytic domain (3A6B4, 1:500), mouse anti-Wingless (Wg) (4D4, 1:500), mouse anti-Nubbin (Nub) (Nub 2D4, 1:25), mouse anti-LacZ (40-1a, 1:100), mouse anti-Achaete (Ac) (1:100), mouse anti-Cut (Ct) (2B10, 1:100), mouse anti-Delta extracellular (C594.9B-s, 1:100), mouse anti-Discs large (Dlg) (4F3, 1:100). Primary antibodies from Cell Signaling include rabbit anti-HA (C29F4, 1:1000), mouse anti-HA (6E2, 1:1000), rabbit anti-cleaved Drosophila Dcp-1 (Asp215, 1:1000), mouse anti-P-histone H3 (Ser10) (PH3) (6G3, 1:1000), and rabbit Phoso-SMAD1/5 (Ser463/465) (41D10, 1:1000). Primary antibodies from Abcam include chicken anti-GFP (ab13970, 1:1000). Generously gifted antibodies include rabbit anti-Vestigial (Vg) (1:500) (gift from Dr. Kirsten Guss), mouse anti-Blistered (Bs) (gift from Dr. Seth Blair), rabbit anti-Salm (Sal) (1:500) (gift from Dr. Baotong Xie and Dr. Brian Gebelein), rat anti-Araucan (Ara/Caup) (1:200) (gift from Dr. Sonsoles Campuzano), rat anti-Aristaless (Al) (1:1000) (gift from Dr. Gerard Campbell). Secondaries (1:500) were obtained from Invitrogen and DAPI (1:500) was used as a nuclear stain. For Wg vesicle staining, a modified protocol was used to enhance signal detection by increasing the secondary antibody concentration. Because proteins in vesicles are commonly present in minimal amounts, this adjustment improves fluorescence intensity by maximizing secondary antibody binding to each primary thereby amplifying the signal. Affinity Photo was used to process all images.

**Asperous Antibody Generation**

Custom rabbit monospecific antibodies against Aspr (NM_134528.3) were generated by Pacific Immunology using three synthetic peptides corresponding to residues 33-49 (CTYRTYYTYGDGRSLQR), 45-67 (RSLQRVVYRDPVYTRAQSYASGC), and 429-441 (CRQTRTFPVAKYH). Each peptide was conjugated to keyhole limpet hemocyanin (KLH) and used to immunize two rabbits over a 13-week protocol. Pre-immune serum was collected prior to immunization, followed by four immunizations and four production bleeds per animal using AdjuliteTM adjuvant. Antibody titers were assessed by ELISA at the first production bleed, and approximately 200 mL of total serum was collected across all bleeds. Serum (~25mL) was affinity-purified against the immunizing peptides using a column, yielding monospecific IgG antibodies.

**Hybridization Chain Reaction RNA Fluorescence in situ Hybridization (HCR RNA FISH)**

HCR RNA-FISH was performed using Molecular Instruments’ HCR™ RNA-FISH kit, with adaptations to the protocol made to accommodate our specific tissue requirements. L3 larvae were inverted in 1× PBS, fixed in 4% PFA for 20 min at 4°C, dehydrated through 50%, 70%, and 100% ethanol (EtOH) washes, and then partially rehydrated in 50% EtOH/0.1% PBST before final washes in 0.1% PBST. Samples were pre-hybridized in Probe Hybridization Buffer for 30 min at 37°C, followed by overnight incubation at 37°C with 2 pmol of each RNA probe in fresh Probe Hybridization Buffer. The next day, unbound probes were removed by washing at 37°C in Probe Wash Buffer, then three washes in 5X Sodium Chloride Sodium Citrate (SSCT) at room temperature. For amplification, tissues were incubated in Amplification Buffer for 30 min in the dark, and HCR hairpins (h1 and h2, 30 pmol each) were snap-cooled separately by heating at 95°C for 90 s. The hairpin mixture was then added, and samples were incubated for 12 h to 16 h at room temperature in the dark. Following five washes in 5X SSCT, samples were stored at 4°C until dissection and imaging. Custom probes were designed for aspr (NM_134528) and eGFP (KC896842).

**Edu Staining**

Edu incorporation using the Click-iT EdU Alexa Fluor Imagina Kit (Invitrogen) was used to assess cell proliferation. Third-instar larvae were inverted in Schneider’s media, then incubated in a solution of 1 µL 10 µM EdU per 1 mL Schneider’s media for 20 min. After removing the EdU solution, carcasses were fixed in 4% PFA for 20 min and washed in 0.1% PBT. IHC, if necessary, was carried out at this point. The Click-iT Reaction mixture (Reaction Buffer, Buffer Additive, CuSO₄, and Alexa Fluor azide) were prepared according to the kit instructions, combined, and immediately added to the samples. Tissues were incubated for 30 min in the dark, followed by washes in PBS containing DAPI or Hoechst 33342 to label nuclei. Samples were stored in PBS at 4°C until dissection and imaging (typically within 48 h).

**Hemolymph Collection and Western Blot Analysis**

To determine the extracellular presence of Aspr, hemolymph was extracted from 30 L3 larvae of W1118 and ci-Gal4 x Aspr-HA genotypes. Larvae were bled in 200 uL of PBS for 10 mins and briefly centrifuged (700 x g, 30s) through a 40µm Mini Strainer (pluriSelect USA) to filter debris. The filtrate and debris were collected separately. The filtrate was further centrifuged at 17,000 x g for 1 h at 4°C to pellet dense material. Both the pellet and the supernatant were resuspended in PBS (100 µL and 200 µL, respectively) containing protease inhibitors (Sigma), treated with 2.5 µL Insect Pop Culture Reagent (Sigma), and incubated for 15 mins at room temperature. Larvae lysates and pellet from the centrifugation were then sonicated using a Branson M28000 water bath sonicated (three to 5 cycles; 5 min on, 2 mins off).

Samples were mixed with SDS-PAGE buffer (10% glycerol, 60nM Trish-HCI pH 6.8, 2% SDS, 0.01 bromophenol blue, 1.25% β-mercaptoethanol), heated at 95°C for 10 mins, and 30 µL was loaded onto 10% SDS-PAGE gels. After electrophoresis, proteins were transferred to 0.45 µm PVDF membranes (Bio-Rad) using the eBlot System (GenScript), blocked in 5% milk in PBST (0.05% Tween-20) for 1 h, and probed with rabbit anti-HA (Cell Signaling, C29F4, 1:2500) or mouse anti-actin (Abcam, 1:5000). After washing, membranes were incubated with HRP-conjugated secondary anti-rabbit or anti-mouse IgG (Jackson ImmunoResearch, 515-035-062, 1:10,000) for 30 mins and developed using Immun-Star Chemiluminescenes (Bio-Rad). Images were acquired using an Amersham ImageQuant 800 imagers (Cytiva).

**Quantification and Statistical Analysis**

**Statistical Tests**

Statistical analyses were performed using GraphPad Prism Software. Statistical significance for all quantified data was determined by ANOVA followed by Bonferroni post-hoc tests, as detailed in each figure legend. Specific sample sizes (n values), statistical tests, and significance criteria (p values) are indicated in figure legends. Data is presented as means ± standard deviation (SD) or ± standard error (SE), as indicated in individual figure legends.

**Regeneration Scoring and Wing Measurements**

Adult flies were anesthetized, and wings were categorized by distinct regenerative phenotypes as described in Figure 2. Regeneration data for each experiment is illustrated as stacked bar graphs, showing the percentage of wings in each scoring group. Wings were then removed, mounted in Fisher Chemical^TM^ Permount^TM^ Mounting Medium (Fischer Scientific) and imaged using a Zeiss Discovery.V8 microscope. Wing area was quantified with Fiji (ImageJ), with males and females analyzed separately to account for sex-specific size differences. To exclude the hinge region in measurements, the L1 vein was traced until encountering the vertically translucent area between the hinge and the wing blade followed by tracing of the L6 vein. Statistical analysis was performed using GraphPad Prism 10.2.

**Developmental Timing Experiments**

Developmental timing was assessed by monitoring pupation and eclosion rates at 18^o^C. Eggs were collected in cages over a 6 hr time window at 22^o^C, after which 50 eggs were transferred into each vial and shifted to 18^o^C for developmental monitoring. Pupation and eclosion were scored every 12 hours. Pupation was scored on the presence of a fully formed light-brown puparium, whereas eclosion was defined as the complete exit of the adult fly from the pupal shell and physical detachment. At each time point, the number of pupae and newly eclosed adults were recorded, with adults sexed upon observation. Genotypes of flies are listed in Sup Fig. 2.

**Fluorescence Intensity Profiling and quantification**

To assess spatial patterns of proliferation activity, fluorescence intensity of PCNA-GFP was quantified. For each disc, a straight line was drawn along the D/V boundary, encompassing from A/P compartments, to capture intensity across the full width of the pouch. Fluorescence values along this line were extracted using the Profile tool in Zeiss ZEN imaging software, which reports pixel intensity values at regular intervals along the line. Profiles were generated by scaling each disc to percent disc width, averaging fluorescence values at equivalent positions across 3-4 discs per time point. A line of best fit was generated using E. Lowe’s method. Data was analyzed using GraphPad Prism. Quantification of *aspr* RNA (Figure 1) was performed on experiments using HCR RNA-FISH (described above) to visualize Aspr transcripts at the indicated time points following ablation. Fluorescence intensity of the pouch was measured using Fiji and normalized to the notum (excluding the *aspr* signal at the tip of the notum). Pouch and notum were identified using disc morphology. Ten discs were used per time point.

**Vesicle size quantification**

Images acquired at 40X magnification were imported into FIJI (ImageJ) and calibrated using a 10 µm scale bar from ZEN Pro. Pixel length was determined from the scale bar (89 pixel = 10 µm), and this calibration was applied globally to all three wing disc images. Each image was then viewed at 150% zoom, and 20 vesicles per disc were measured.

**Computational Analysis**

Structural predictions of full-length Aspr and the signal peptide regions of aspr, Dumpy, NELL2 (human), and EDIL3 (mouse and human) were generated using AlphaFold v3. Predicted structures were visualized and prepared using PyMOL (Schrödinger, LLC). Predicted Aligned Error (PAE) for full-length Aspr and for each signal peptide structure was computed using UCSF ChimeraX 1.9 to evaluate the confidence in residue positioning and domain orientation.

To predict the presence of a signal peptide and infer subcellular localization of Aspr, six publicly available prediction tools were used: SignalP, Phobius, TargetP, Protter, TMHMM-2.0, and DeepLoc. Prediction scores from each tool were compiled and visualized as a heat map using Python. Putative homologs of aspr were identified using sequence searches on FlyBase and the DALI structure comparison server. Signal peptide sequences for Dumpy, EDIL3 (mouse and human), and NELL2 were retrieved from UniProt. Sequences were aligned using Clustal Omega and visualized with Jalview 2.11.4.1. Signal peptide structures were predicted with AlphaFold v3, and pairwise structural alignments were performed in PyMOL to calculate root-mean-square deviation (RMSD) values.

**Figure Illustration and Visualization Tools**

Schematics and conceptual diagrams were created using BioRender (BioRender.com). BioRender was used to generate schematics for the a*spr* DRMS (Sup Fig. 1I), gene regulatory network (Sup Fig. 6F), and Aspr functional model (Fig. 9). All illustrations were produced under an academic license and are intended for publication and educational use.
