## Supplemental Figures for "Asperous coordinates regenerative timing by regulating damage-induced WNT Signaling"

### Figure S1

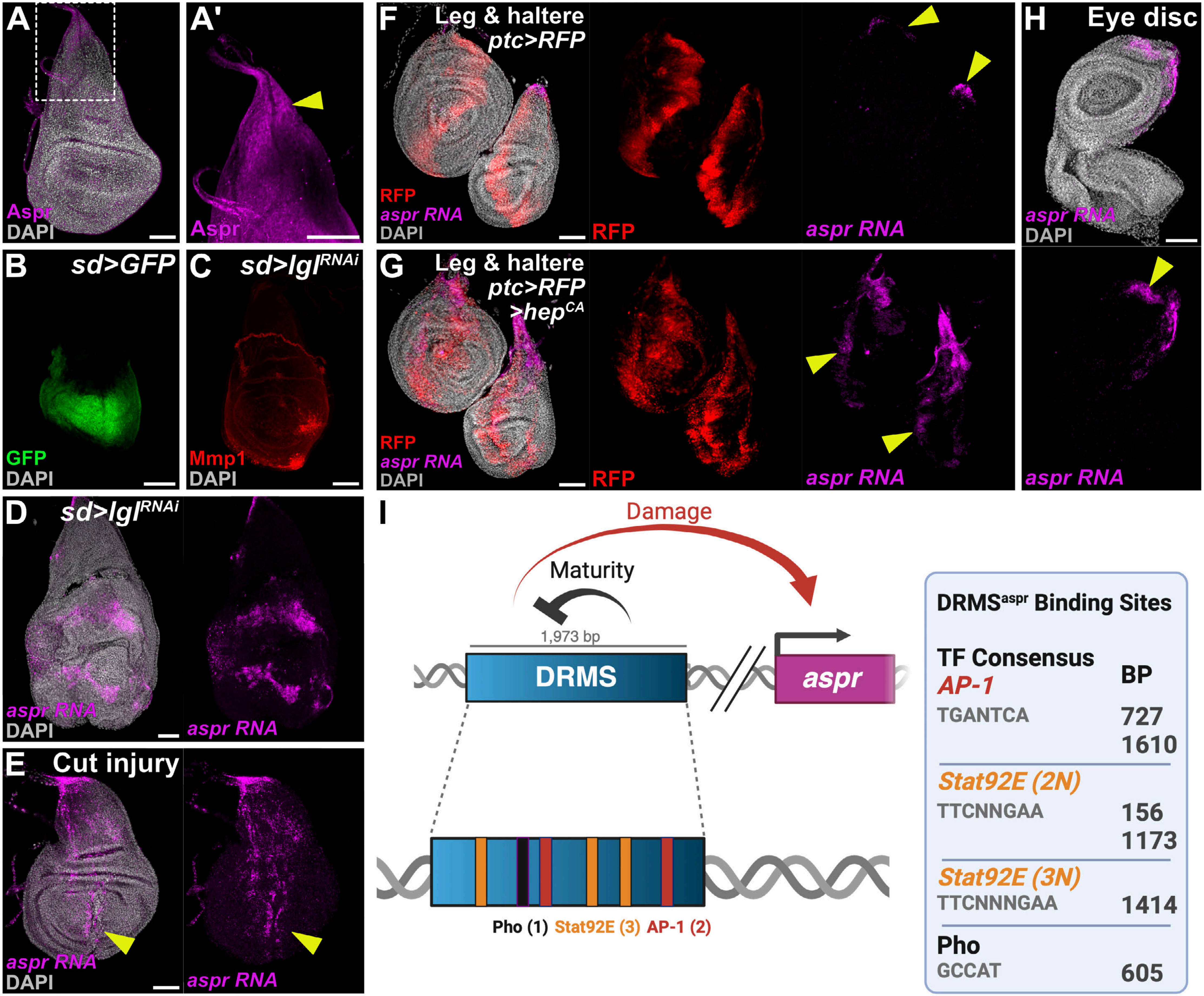

A
Aspr
DAPI
A'
Aspr
B
sd>GFP
GFP
DAPI
C
sd>lgl^RNAi
Mmp1
DAPI
D
sd>lgl^RNAi
aspr RNA
DAPI
aspr RNA
E
Cut injury
aspr RNA
DAPI
aspr RNA
F
Leg & haltere
ptc>RFP
RFP
aspr RNA
DAPI
RFP
aspr RNA
G
Leg & haltere
ptc>RFP
>hep^CA
RFP
aspr RNA
DAPI
RFP
aspr RNA
H
Eye disc
aspr RNA
DAPI
aspr RNA
I
Damage
Maturity
1,973 bp
DRMS
aspr
Pho (1) Stat92E (3) AP-1 (2)
DRMS^aspr Binding Sites
TF Consensus
BP
AP-1
TGANTCA
727
1610
Stat92E (2N)
TTCNNGAA
156
1173
Stat92E (3N)
TTCNNNGAA
1414
Pho
GCCAT
605

**Supplemental Figure Legends**

**Figure S1. *aspr* Expression in Imaginal Discs – related to Figure 1**

(**A-A'**) Wild type $w^{1118}$ wing disc stained with anti-Aspr antibody (magenta) shows developmental expression in the notum. DAPI (grey) marks nuclei.

(**B**) Early L3 disc expressing *UAS-GFP* under *sd-GAL4* highlights the endogenous *sd* expression domain.

(**C**) *sd-GAL4>lgl*$^{RNAi}$ early L3 discs showing neoplastic tumors. Mmp1 (red) marks regions of JNK activity within the *sd* domain.

(**D**) *sd-GAL4>lgl*$^{RNAi}$ strongly upregulates *aspr* RNA (magenta) in response to tumor formation.

(**E**) Physically damaged wild type early L3 disc shows *aspr* RNA upregulation near the injury site (yellow arrowhead).

(**F**) Control *ptc-GAL4>UAS-RFP* early L3 leg and haltere discs show RFP (red) along the A/P boundary and developmental *aspr* RNA in the dorsal thorax of the haltere and dorsal leg (yellow arrowheads).

(**G**) *ptc-GAL4>UAS-RFP,UAS-hepCA* early L3 discs show robust induction of *aspr* RNA across the tissue (yellow arrowheads) in response to damage signaling.

(**H**) Wild type $w^{1118}$ L3 eye disc shows developmental *aspr* RNA in the antennal field.

(**I**) Schematic of the Damage-Responsive Maturity-Silenced (DRMS) enhancer of *aspr*, including a zoomed-in view of regulatory binding sites (Pho, Stat92E, AP-1), with genomic positions listed in the accompanying table. Scale bars: 50 µm unless indicated. Full genotypes are provided in Supplementary Genotypes.

### Figure S2

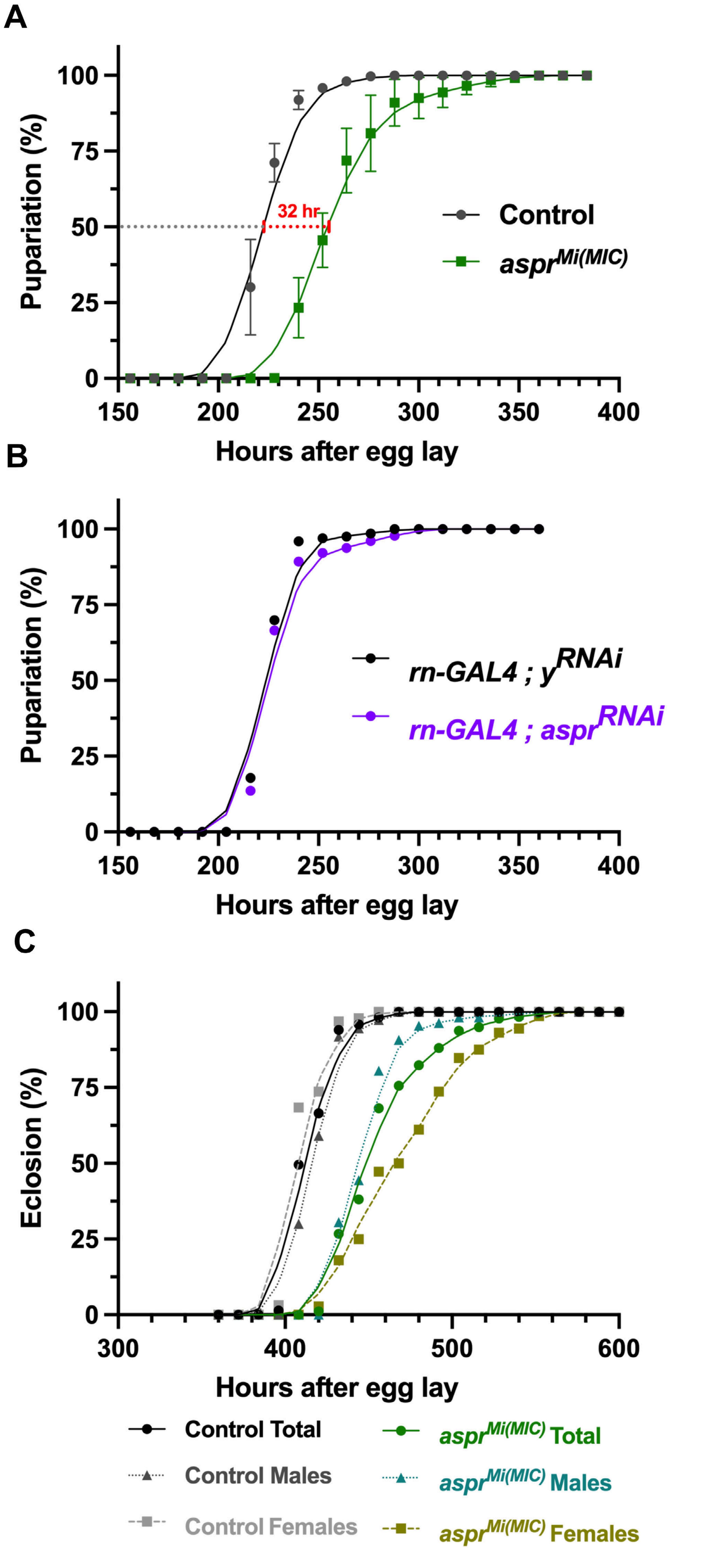
A
Pupariation (%)
100
75
50
25
0
150
200
250
300
350
400
Hours after egg lay
32 hr
Control
aspr^Mi(MIC)
B
Pupariation (%)
100
75
50
25
0
150
200
250
300
350
400
Hours after egg lay
rn-GAL4 ; y^RNAi
rn-GAL4 ; aspr^RNAi
C
Eclosion (%)
100
75
50
25
0
300
400
500
600
Hours after egg lay
Control Total
Control Males
Control Females
aspr^Mi(MIC) Total
aspr^Mi(MIC) Males
aspr^Mi(MIC) Females

**Figure S2. Loss of *aspr* Delays Development and Eclosion – related to Figure 2**

(**A**) Pupation timing of $aspr^{Mi(MIC)}$ (green squares) compared with wild type controls (grey circles). X-axis, % pupated; Y-axis, hours after egg lay. $aspr^{Mi(MIC)}$ larvae reach 50% pupation ~32 h later than controls, indicating delayed development. Mean ± SD; 3 biological replicates. Sample sizes: Wild type $w^{1118}$ control (n = 157, 233, 290); $aspr^{Mi(MIC)}$ (n = 122, 202, 226).

(**B**) Pupation timing of *rn-GAL4>*$aspr^{RNAi}$ (purple circles) versus *rn-GAL4>*$y^{RNAi}$ controls (black circles). Reducing *aspr* in the pouch does not alter overall developmental timing. Sample sizes: *rn-GAL4>*$y^{RNAi}$ (n = 196); *rn-GAL4>*$aspr^{RNAi}$ (n = 176).

(**C**) Eclosion timing of $aspr^{Mi(MIC)}$ and wild type $w^{1118}$, separated by sex. Control females eclose earlier than males, whereas the opposite pattern occurs in $aspr^{Mi(MIC)}$, where hemizygous null males eclose earlier than homozygous null females. $aspr^{Mi(MIC)}$ also shows a reduced overall eclosion rate. Sample sizes: $w^{1118}$ males (n = 95), females (n = 110); $aspr^{Mi(MIC)}$ males (n = 72), females (n = 108).

Full genotypes are provided in Supplementary Genotypes.

### Figure S3

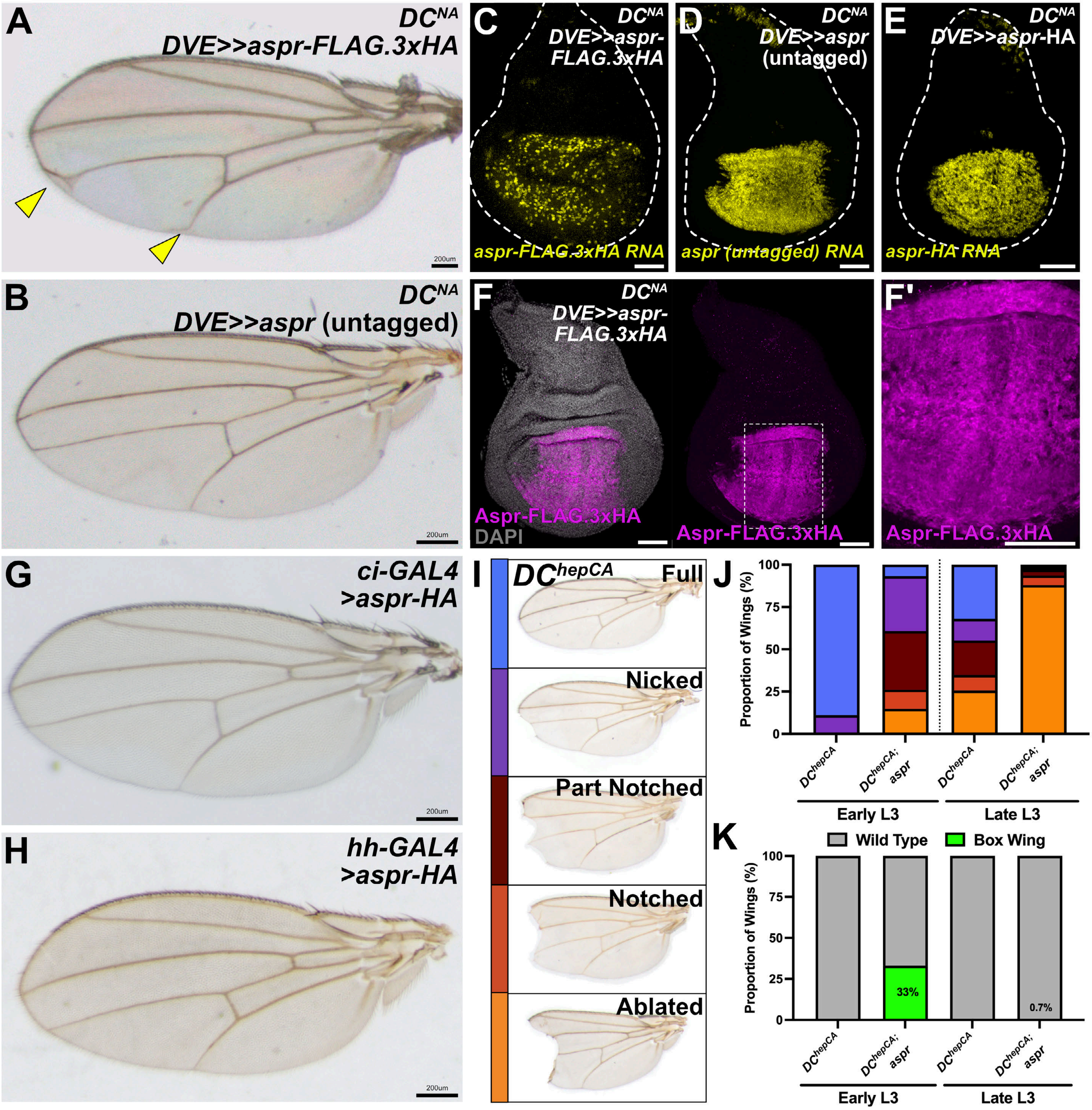

A
DC^NA
DVE>>aspr-FLAG.3xHA
200um
B
DC^NA
DVE>>aspr (untagged)
200um
C
DC^NA
DVE>>aspr-FLAG.3xHA
aspr-FLAG.3xHA RNA
D
DC^NA
DVE>>aspr (untagged)
aspr (untagged) RNA
E
DC^NA
DVE>>aspr-HA
aspr-HA RNA
F
DC^NA
DVE>>aspr-FLAG.3xHA
Aspr-FLAG.3xHA
DAPI
Aspr-FLAG.3xHA
F'
Aspr-FLAG.3xHA
G
ci-GAL4 >aspr-HA
200um
H
hh-GAL4 >aspr-HA
200um
I
DC^hepCA
Full
Nicked
Part Notched
Notched
Ablated
J
Proportion of Wings (%)
100
75
50
25
0
DC^hepCA
DC^hepCA; aspr
DC^hepCA
DC^hepCA; aspr
Early L3
Late L3
K
Wild Type
Box Wing
Proportion of Wings (%)
100
75
50
25
0
33%
0.7%
DC^hepCA
DC^hepCA; aspr
DC^hepCA
DC^hepCA; aspr
Early L3
Late L3

**Figure S3. Developmental and Regenerative Effects of Distinct *Aspr* Constructs – related to Figure 3**

(**A–B**) Adult wings from unablated early L3 discs (*DC*$^{NA}$*, DVE>>GAL4*) ectopically expressing *aspr-FLAG.3xHA* or untagged *aspr*. Aspr-FLAG.3xHA (A) causes vein detachments from the posterior margin (yellow arrowheads), whereas untagged Aspr (B) does not alter wing morphology.

(**C–E**) Unablated early L3 discs (*DC*$^{NA}$*, DVE>>GAL4*) expressing *aspr* constructs and probed for *aspr* RNA (yellow). Untagged *aspr* (D) and *aspr*-HA (E) show uniform RNA expression, whereas *aspr-FLAG.3xHA* (C) produces patchy signal, suggesting RNA instability or degradation.

(**F–F'**) Aspr-FLAG.3xHA protein (magenta) in unablated discs (*DC*$^{NA}$*, DVE>>GAL4*) shows uniform distribution but lacks the punctate accumulation characteristic of Aspr-HA (main Figure 3A',B'). DAPI (grey) marks nuclei. Dashed outline in (F) shows magnified area in (F').

(**G–H**) Adult wings from unablated early L3 discs (*DC*$^{NA}$) expressing *aspr-HA* in the anterior compartment via *ci-GAL4* (G) or posterior compartment via *hh-GAL4* (H) shows no developmental wing defects.

(**I**) Adult wing phenotype categories used for regeneration scoring from ablated wing discs (*DC*$^{hepCA}$).

(**J**) Regeneration scoring of adult wings from early and late L3 ablated discs (*DC*$^{hepCA}$*, DVE>>GAL4*) expressing untagged *aspr*. Proportions of wings in each outcome class are plotted. Ectopic Aspr reduces regenerative capacity at both stages, similar to Aspr-HA (Main Figure 3M).

(**K**) Proportion of wings exhibiting the box-wing phenotype (green) vs. wild-type regenerative outcomes (grey) in the experiment in (J). Box-wing defects occur in 33% of early L3 ablations but only 0.7% of late L3 ablations. Sample sizes listed in text.

Scale bars: 50 µm unless indicated. Full genotypes in Supplementary Genotypes.

### Figure S4

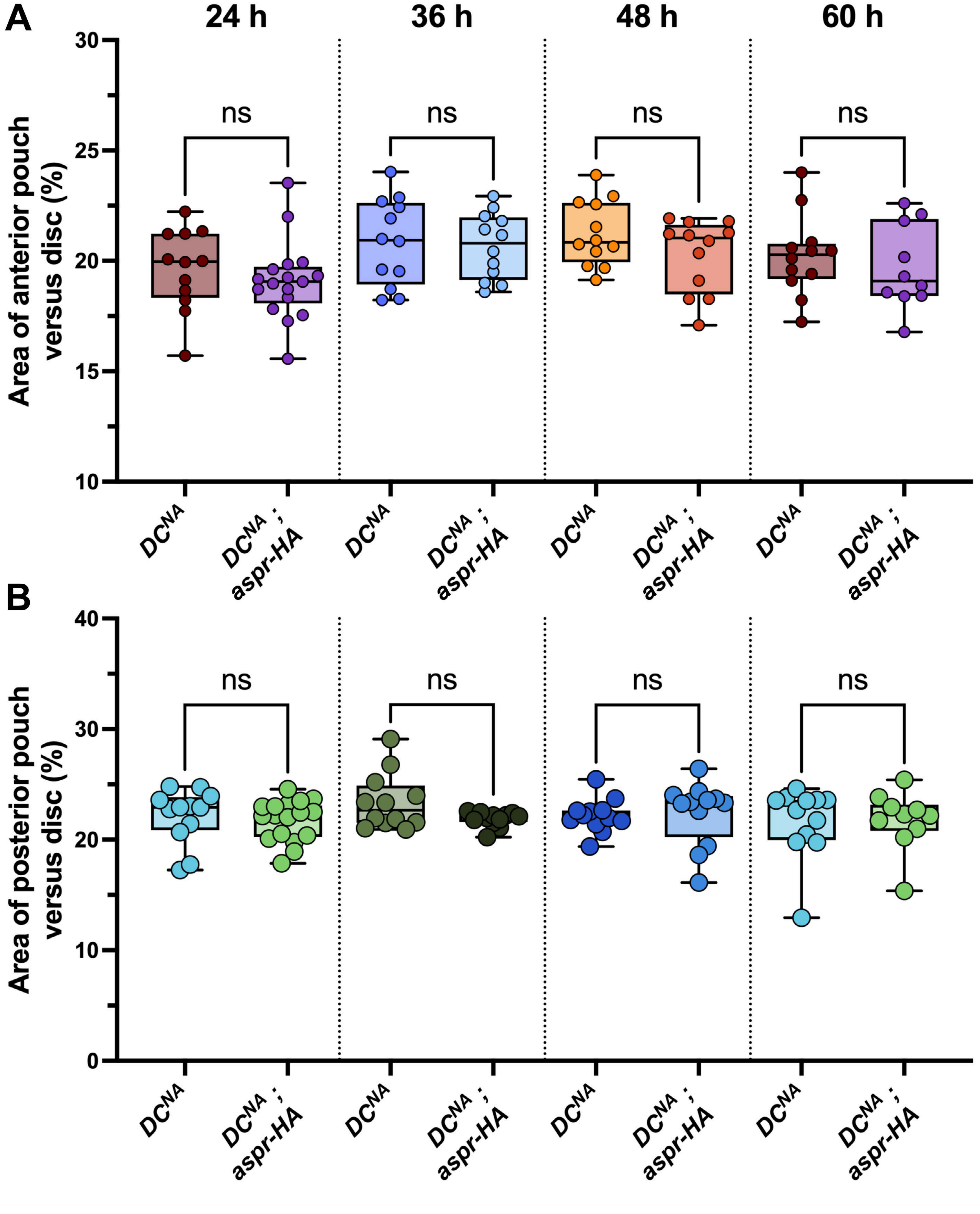

A
24 h
36 h
48 h
60 h
ns
ns
ns
ns
Area of anterior pouch versus disc (%)
30
25
20
15
10
DCNA
DCNA ; aspr-HA
DCNA
DCNA ; aspr-HA
DCNA
DCNA ; aspr-HA
DCNA
DCNA ; aspr-HA
B
ns
ns
ns
ns
Area of posterior pouch versus disc (%)
40
30
20
10
0
DCNA
DCNA ; aspr-HA
DCNA
DCNA ; aspr-HA
DCNA
DCNA ; aspr-HA
DCNA
DCNA ; aspr-HA

**Figure S4. Ectopic *Aspr* Does Not Alter Pouch Growth – related to Figure 4**

(**A–B**) Quantification of anterior (A) and posterior (B) pouch areas from unablated discs (*$DC^{NA}$, DVE>>GAL4*) (Main Figure 3G) expressing *$y^{RNAi}$* (control) or *Aspr*-HA from 24–60 h post-ablation. Measurements are shown as percent of total disc area. No significant differences were observed at any time point (one-way ANOVA). Box plots show median, quartiles, and data distribution. Ns = not significant. Sample sizes listed in text.

### Figure S5

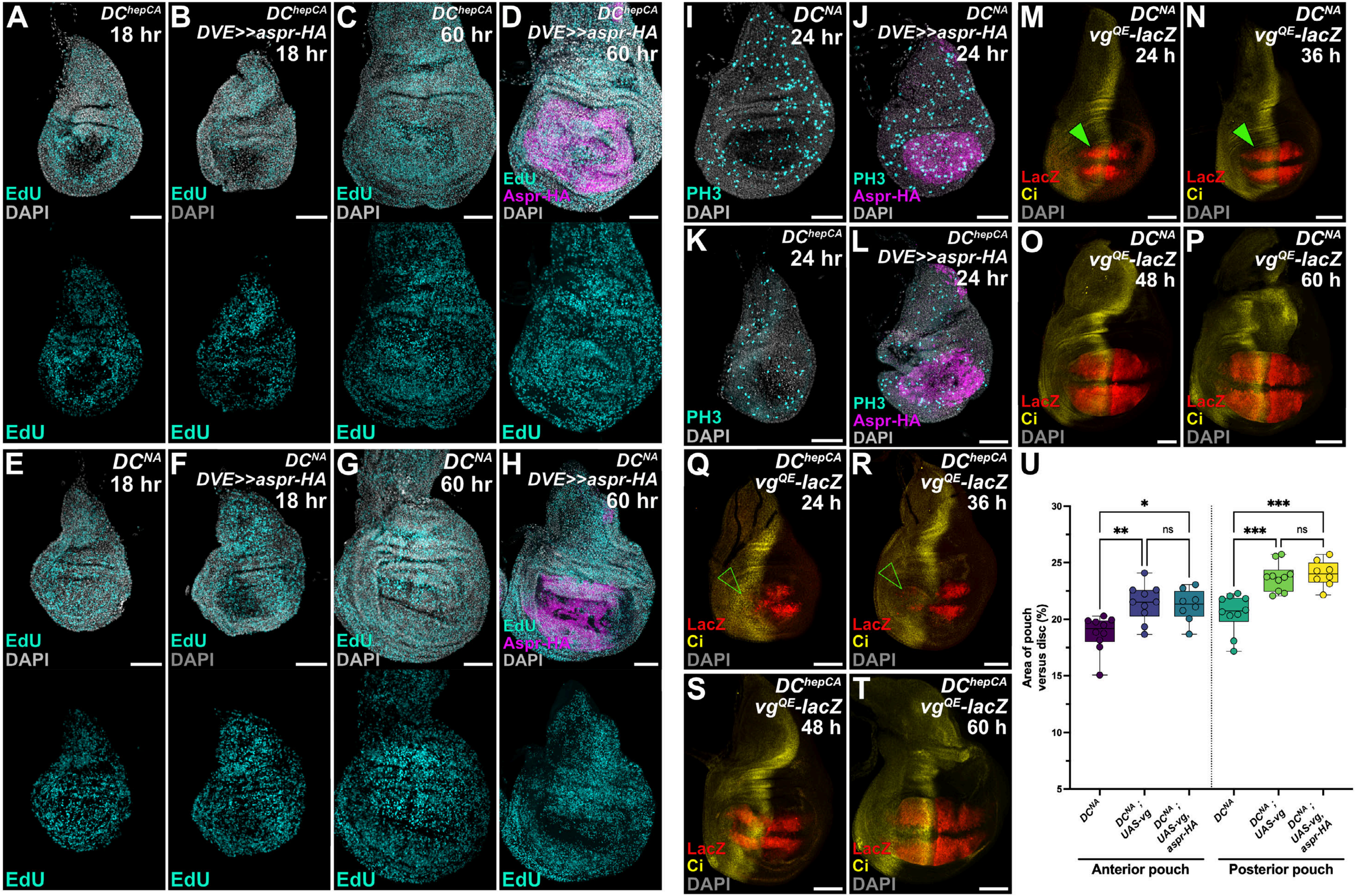
A
$DC^{hepCA}$
18 hr
EdU
DAPI
EdU
B
$DC^{hepCA}$
DVE>>aspr-HA
18 hr
EdU
DAPI
EdU
C
$DC^{hepCA}$
60 hr
EdU
DAPI
EdU
D
$DC^{hepCA}$
DVE>>aspr-HA
60 hr
EdU
Aspr-HA
DAPI
EdU
E
$DC^{NA}$
18 hr
EdU
DAPI
EdU
F
$DC^{NA}$
DVE>>aspr-HA
18 hr
EdU
DAPI
EdU
G
$DC^{NA}$
60 hr
EdU
DAPI
EdU
H
$DC^{NA}$
DVE>>aspr-HA
60 hr
EdU
Aspr-HA
DAPI
EdU
I
$DC^{NA}$
24 hr
PH3
DAPI
J
$DC^{NA}$
DVE>>aspr-HA
24 hr
PH3
Aspr-HA
DAPI
K
$DC^{hepCA}$
24 hr
PH3
DAPI
L
$DC^{hepCA}$
DVE>>aspr-HA
24 hr
PH3
Aspr-HA
DAPI
M
$DC^{NA}$
$vg^{QE}$-lacZ
24 h
LacZ
Ci
DAPI
N
$DC^{NA}$
$vg^{QE}$-lacZ
36 h
LacZ
Ci
DAPI
O
$DC^{NA}$
$vg^{QE}$-lacZ
48 h
LacZ
Ci
DAPI
P
$DC^{NA}$
$vg^{QE}$-lacZ
60 h
LacZ
Ci
DAPI
Q
$DC^{hepCA}$
$vg^{QE}$-lacZ
24 h
LacZ
Ci
DAPI
R
$DC^{hepCA}$
$vg^{QE}$-lacZ
36 h
LacZ
Ci
DAPI
S
$DC^{hepCA}$
$vg^{QE}$-lacZ
48 h
LacZ
Ci
DAPI
T
$DC^{hepCA}$
$vg^{QE}$-lacZ
60 h
LacZ
Ci
DAPI
U
Area of pouch versus disc (%)
30
25
20
15
10
5
*
**
ns
***
***
ns
$DC^{NA}$
$DC^{NA}$ ; UAS-vg
$DC^{NA}$ ; UAS-vg, aspr-HA
$DC^{NA}$
$DC^{NA}$ ; UAS-vg
$DC^{NA}$ ; UAS-vg, aspr-HA
Anterior pouch
Posterior pouch

**Figure S5. Ectopic *Aspr* Does Not Affect EdU or PH3 Proliferation Markers – related to Figure 5**

(**A–H**) Ablated discs ($DC^{hepCA}$, A–D) and unablated discs ($DC^{NA}$, E–H) expressing $y^{RNAi}$ (control) or *aspr*-HA with *DVE>>GAL4* at 18 h (A, B, E, F) and 60 h (C, D, G, H) post-ablation stained for EdU incorporation (cyan). DAPI (grey) marks nuclei. No detectable differences in EdU signal were observed.

(**I–L**) Ablated discs ($DC^{hepCA}$, K–L) and unablated discs ($DC^{NA}$, I–J) at 24 h post-ablation stained for PH3 (cyan). Ectopic *aspr*-HA with *DVE>>GAL4* (J, L) does not alter PH3 levels during development or regeneration.

(**M–T**) Time course (24–60 h) of ablated discs ($DC^{hepCA}$, Q–T) and unablated discs ($DC^{NA}$, M–P) bearing $vg^{QE}$-*lacZ* stained for LacZ (red) and Ci (yellow) to mark the A/P boundary. (M–P) Enhancer activity is maintained and expands during development. (Q–T) Following ablation, anterior enhancer activity is lost at 24–36 h (open arrowheads) but reactivated by 48–60 h.

(**U**) Quantification of pouch compartments for non-ablated discs ($DC^{NA}$) in Main Figure 5I–N. Anterior and posterior pouch areas (as % of total disc) are not affected by Aspr-HA. Sample sizes: anterior — control $y^{RNAi}$ n = 10, *vg* n = 10, *vg+aspr-HA* n = 8 ; posterior — control $y^{RNAi}$ n = 10, *vg* n = 10, *vg+aspr-HA* n = 8. Statistics: one-way ANOVA. Annotated p-values: anterior * $p < 0.0119$; posterior ** $p = 0.0024$, posterior *** $p = 0.0002$ ($y^{RNAi}$ to *vg*), *** $p < 0.0001$ ($y^{RNAi}$ to *vg+aspr-HA*); ns = not significant.

Scale bars: 50 µm unless indicated. Full genotypes in Supplementary Genotypes.

### Figure S6

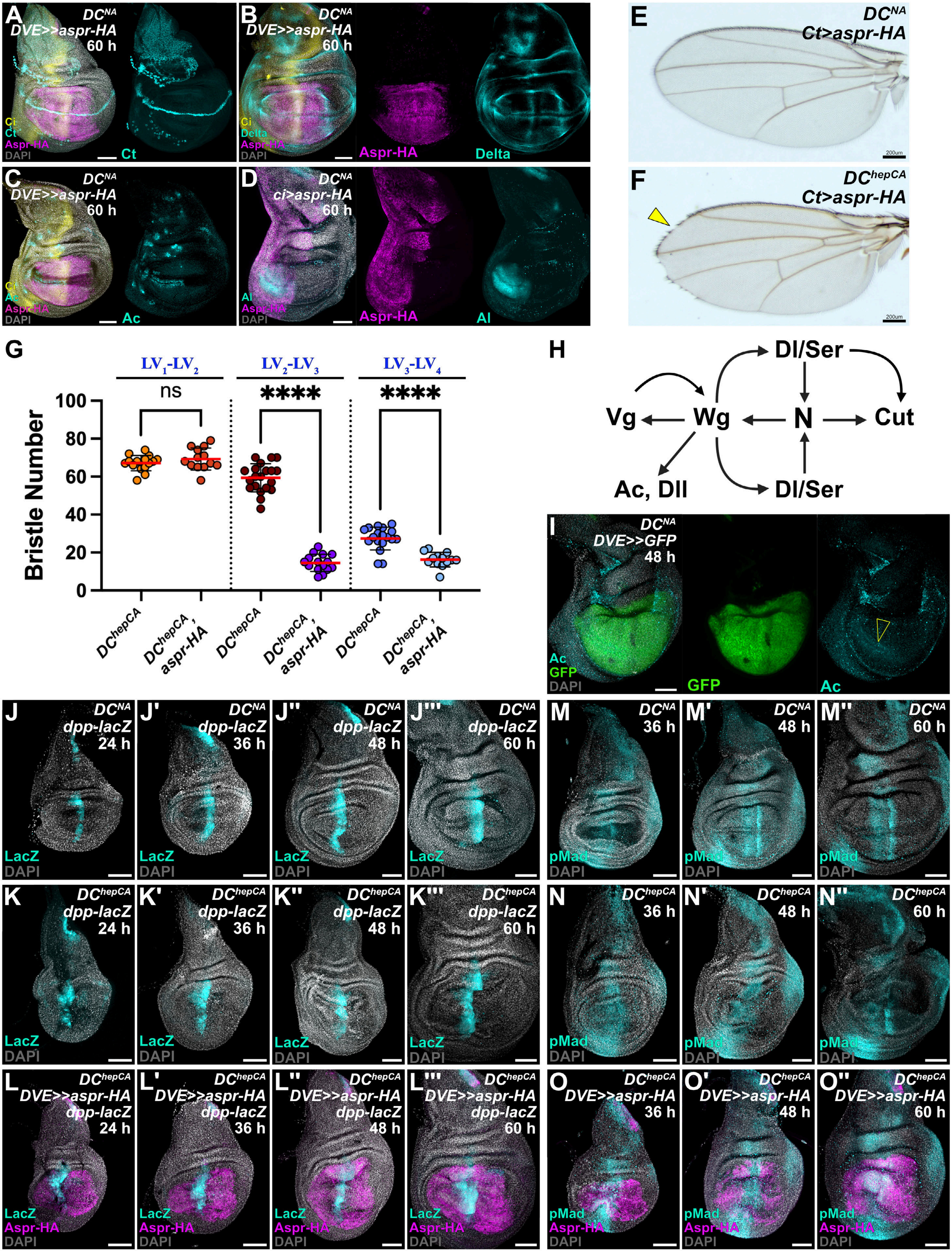
A
DC^NA
DVE>>aspr-HA
60 h
Ci
Ct
Aspr-HA
DAPI
Ct
B
DVE>>aspr-HA
Delta
Aspr-HA
C
Ac
D
ci>aspr-HA
Al
E
Ct>aspr-HA
200um
F
DC^hepCA
G
LV1-LV2
LV2-LV3
LV3-LV4
ns
****
Bristle Number
0
20
40
60
80
100
DC^hepCA
DC^hepCA, aspr-HA
H
Dl/Ser
Vg
Wg
N
Cut
Ac, Dll
I
DVE>>GFP
48 h
GFP
J
dpp-lacZ
24 h
36 h
LacZ
K
L
M
pMad
N
O

**Figure S6. Ectopic *Aspr* Disrupts Anterior Patterning without Altering Dpp – related to Figure 6**

(**A–C**) Unablated ($DC^{NA}$) discs at 60 h expressing *aspr-HA* with *DVE>>GAL4* and stained for Cut (Ct), Delta (Dl), or Achaete (Ac) (cyan). No developmental fate changes are observed.

(**D**) *ci-GAL4>aspr-HA* discs stained for Aristaless (*Al*, cyan) show unaffected *Al* expression.

(**E–F**) Representative adult wings from unablated discs ($DC^{NA}$, E) or ablated discs ($DC^{hepCA}$, F) expressing *aspr-HA* with *ct-GAL4*, showing damage-specific loss of bristles at the margin caused by Aspr in the *ct* domain (yellow arrowhead).

(**G**) Anterior bristle counts along longitudinal veins in wings from early L3 ablations ($DC^{hepCA}$). Aspr-HA significantly reduces bristle number between $LV_2$–$LV_3$ and $LV_3$–$LV_4$; no effect occurs between $LV_1$–$LV_2$. Sample sizes: $LV_1$-LV2: control $y^{RNAi}$ n = 15, *aspr-HA* n = 13 ; $LV_2$–$LV_3$: control $y^{RNAi}$ n = 20, *aspr-HA* n = 15. $LV_3$–$LV_4$: control $y^{RNAi}$ n = 18, *aspr-HA* n = 13 ; Statistics: one-way ANOVA. Annotated p-values: **** $p < 0.0001$; ns = not significant.

(**H)** Schematic of regulatory interactions at the wing margin.

(**I**) Unablated ($DC^{NA}$) disc 48 h post-ablation expressing *GFP* with *DVE>>GAL4* and stained for Achete (Ac, cyan). Achete is not expressed at this developmental stage.

(**G–L'''**) *dpp-lacZ* reporter activity in unablated ($DC^{NA}$) and ablated ($DC^{hepCA}$) discs expressing *aspr-HA* with *DVE>>GAL4* at 24–60 h AHS. Aspr does not affect reporter activity.

(**M–O''**) Discs as in (J–L'''), but at 36 h-60 h and stained for pMad to show Dpp signaling. Aspr does not affect signaling indicated by pMad activity.

Scale bars: 50 μm unless indicated. Full genotypes in Supplementary Genotypes.

### Figure 7 Supplemental

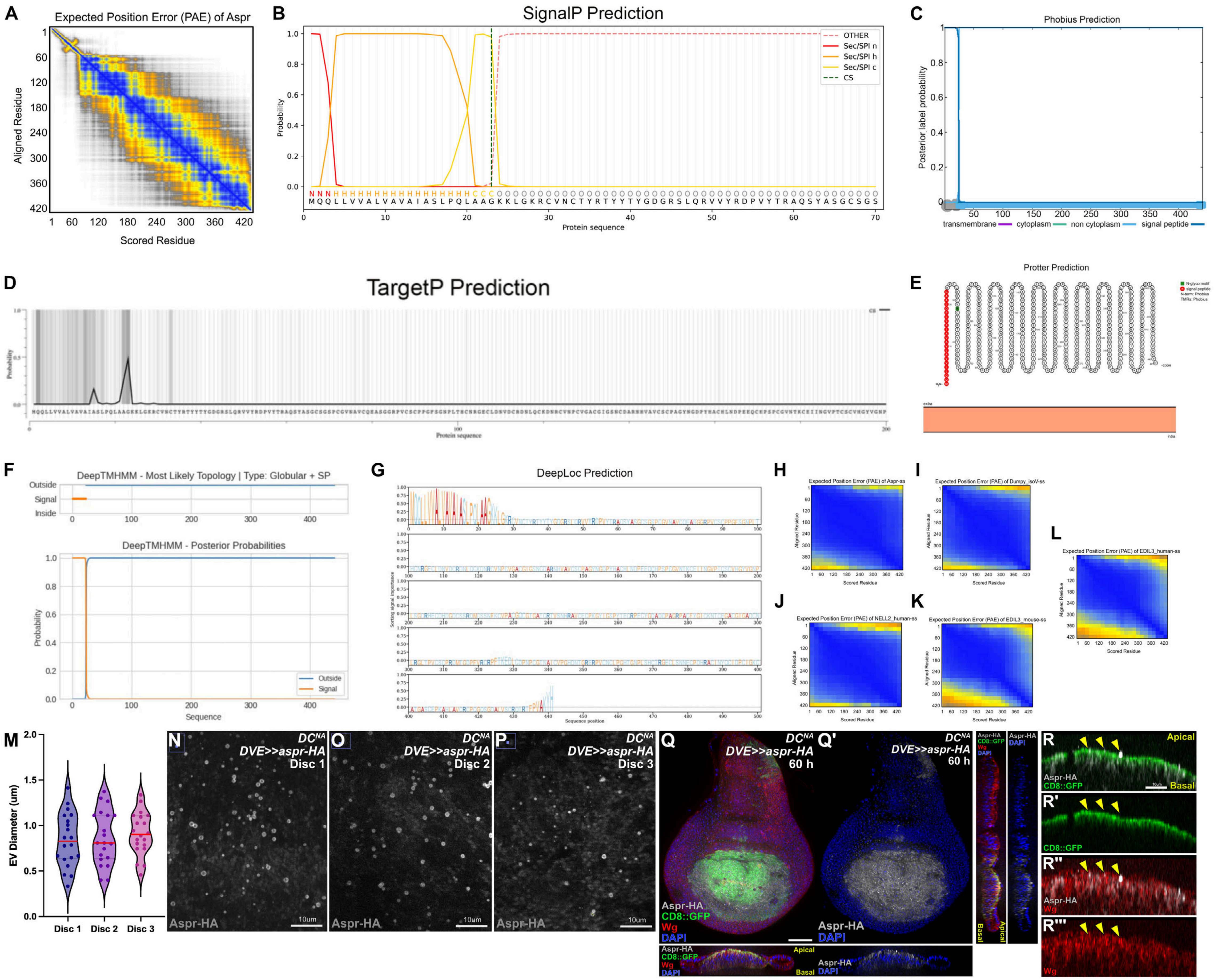

**Figure S7. Structural and Localization Predictions for *Aspr* – related to Figure 7**

(**A**) AlphaFold predicted aligned error (PAE) plot for full-length *Aspr* showing low PAE along the diagonal, consistent with an elongated solenoid architecture with repeated EGF-like domains.

(**B**) SignalP prediction indicating a high-confidence N-terminal signal peptide with a cleavage site between Gly23 and Lys24.

(**C**) Phobius prediction identifies a cleavable signal peptide (residues 1–23) and an extracellular topology with no transmembrane helices.

(**D**) TargetP prediction shows moderate probability (~0.5) for a signal peptide, with no evidence for mitochondrial, chloroplast, or other targeting signals, supporting secretion.

(**E**) Protter visualization of an N-terminal signal sequence (red) and extracellular orientation (orange).

(**F**) DeepTMHMM prediction of a high-confidence signal peptide (residues 1–23) and extracellular localization.

(**G**) DeepLoc identifies *Aspr* as extracellular with high confidence.

(**H–L**) PAE plots for signal peptides of *Aspr* and homologous proteins (Dumpy isoform V, human NELL2, mouse EDIL3, human EDIL3). All show low PAE (blue), indicating confident structural predictions.

(**M**) Quantification of extracellular vesicle (EV) size in three representative discs. n = 20 per disc

(**N-P**) High magnification images of unablated ($DC^{NA}$) discs expressing *aspr-HA* with *DVE>>GAL4* used for EV quantification. Aspr-HA in grey.

(**Q-Q'**) Unablated ($DC^{NA}$) discs expressing *aspr-HA* with *DVE>>GAL4* and expressing membrane *GFP* (*CD8::GFP*) plus aspr-HA (gray) and Wg (red), showing Aspr overlaps with CD8::GFP within cells, but EVs do not have the GFP label.

(**R-R'''**) High-magnification transverse sections of disc in (Q-Q') showing EVs do not colocalize with GFP labelling (yellow arrowhead).

### Figure S8

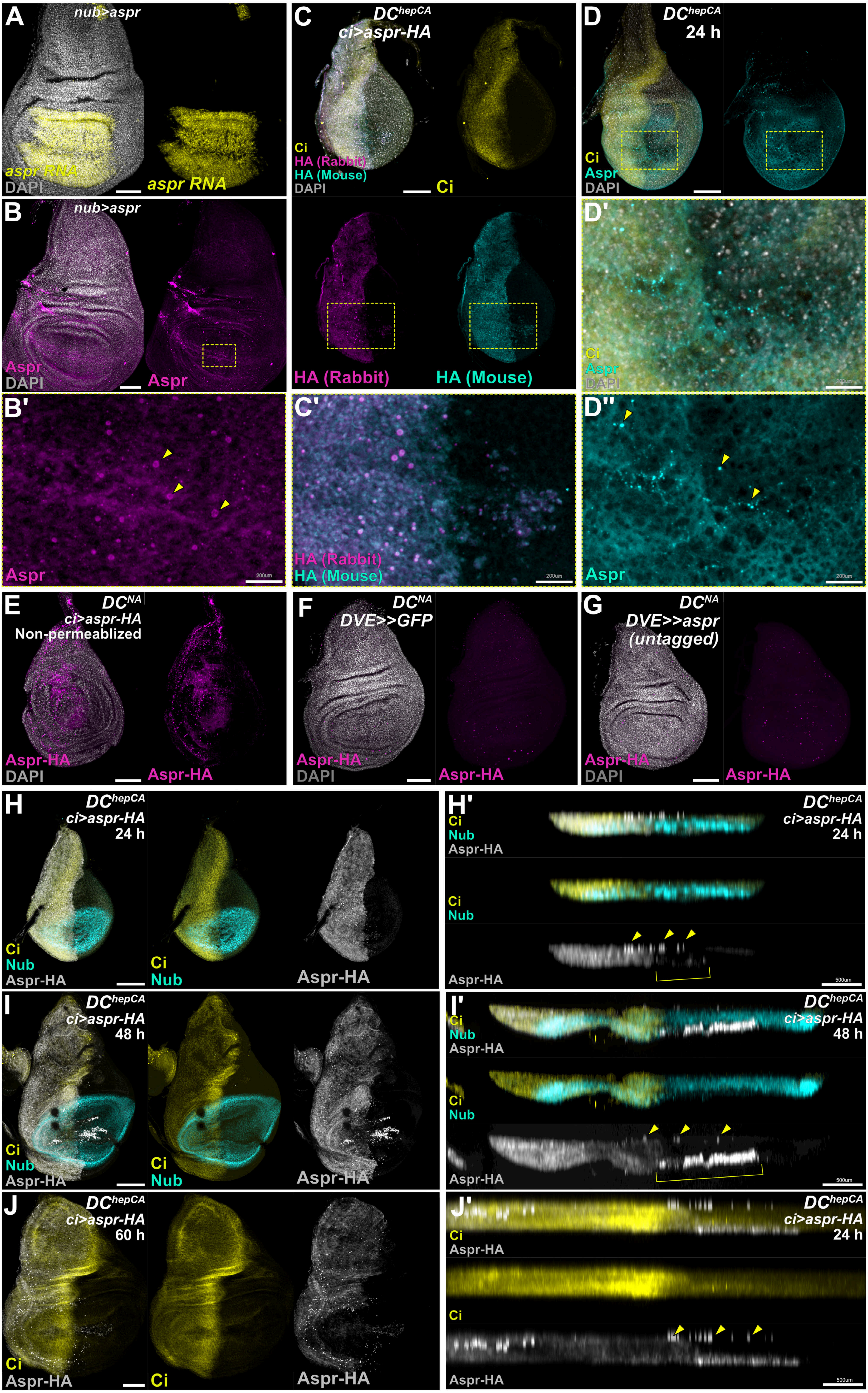
A
nub>aspr
aspr RNA
DAPI
aspr RNA
B
nub>aspr
Aspr
DAPI
Aspr
B'
Aspr
200um
C
DC^hepCA
ci>aspr-HA
Ci
HA (Rabbit)
HA (Mouse)
DAPI
Ci
HA (Rabbit)
HA (Mouse)
C'
HA (Rabbit)
HA (Mouse)
200um
D
DC^hepCA
24 h
Ci
Aspr
DAPI
D'
Ci
Aspr
DAPI
D''
Aspr
200um
E
DC^NA
ci>aspr-HA
Non-permeablized
Aspr-HA
DAPI
Aspr-HA
F
DC^NA
DVE>>GFP
Aspr-HA
DAPI
Aspr-HA
G
DC^NA
DVE>>aspr
(untagged)
Aspr-HA
DAPI
Aspr-HA
H
DC^hepCA
ci>aspr-HA
24 h
Ci
Nub
Aspr-HA
H'
500um
I
DC^hepCA
ci>aspr-HA
48 h
I'
J
DC^hepCA
ci>aspr-HA
60 h
Ci
Aspr-HA
J'
DC^hepCA
ci>aspr-HA
24 h

**Figure S8. Detection of *Aspr* Vesicles – related to Figure 7**

(**A–B')** *nub-GAL4>aspr* (untagged) disc probed for *aspr* RNA (yellow) and stained with *Aspr* antibody (magenta). Zoom-in (B') reveals small *Aspr*-positive vesicles (yellow arrowheads).

(**C–C'**) Ablated disc ($DC^{hepCA}$) expressing *aspr*-HA under *ci-GAL4* stained with mouse anti-HA (magenta) and rabbit anti-HA (cyan) demonstrate detection of *Aspr*-HA vesicles by both antibodies. Ci (yellow) marks the anterior.

(**D–D''**) Unablated disc ($DC^{NA}$) at 24 h AHS stained for endogenous *Aspr* (cyan). Ci (yellow) marks the anterior. Zoom-in shows *Aspr*-positive vesicles (yellow arrowheads).

(**E**) Unablated leg disc ($DC^{NA}$) expressing *aspr*-HA without permeabilization shows extracellular Aspr-HA vesicles.

(**F–G**) Discs expressing GFP or untagged *aspr* stained with anti-HA confirm that HA staining does not generate puncta or EVs, and that untagged *Aspr* is not detectable by HA antibody.

(**H–J'**) Surface views and cross-sections of ablated discs ($DC^{hepCA}$) expressing *aspr-HA* (grey) under *ci-GAL4* at 24, 48, and 60 h post-ablation. Ci marks the anterior and Nub (cyan) marks the pouch. Cross-sections reveal apical Aspr-HA vesicles (yellow arrowheads). Basal debris first appears at 48 h and diminishes by 60 h. Scale bars: 50 µm unless indicated. Full genotypes in Supplementary Genotypes.

### Figure S9

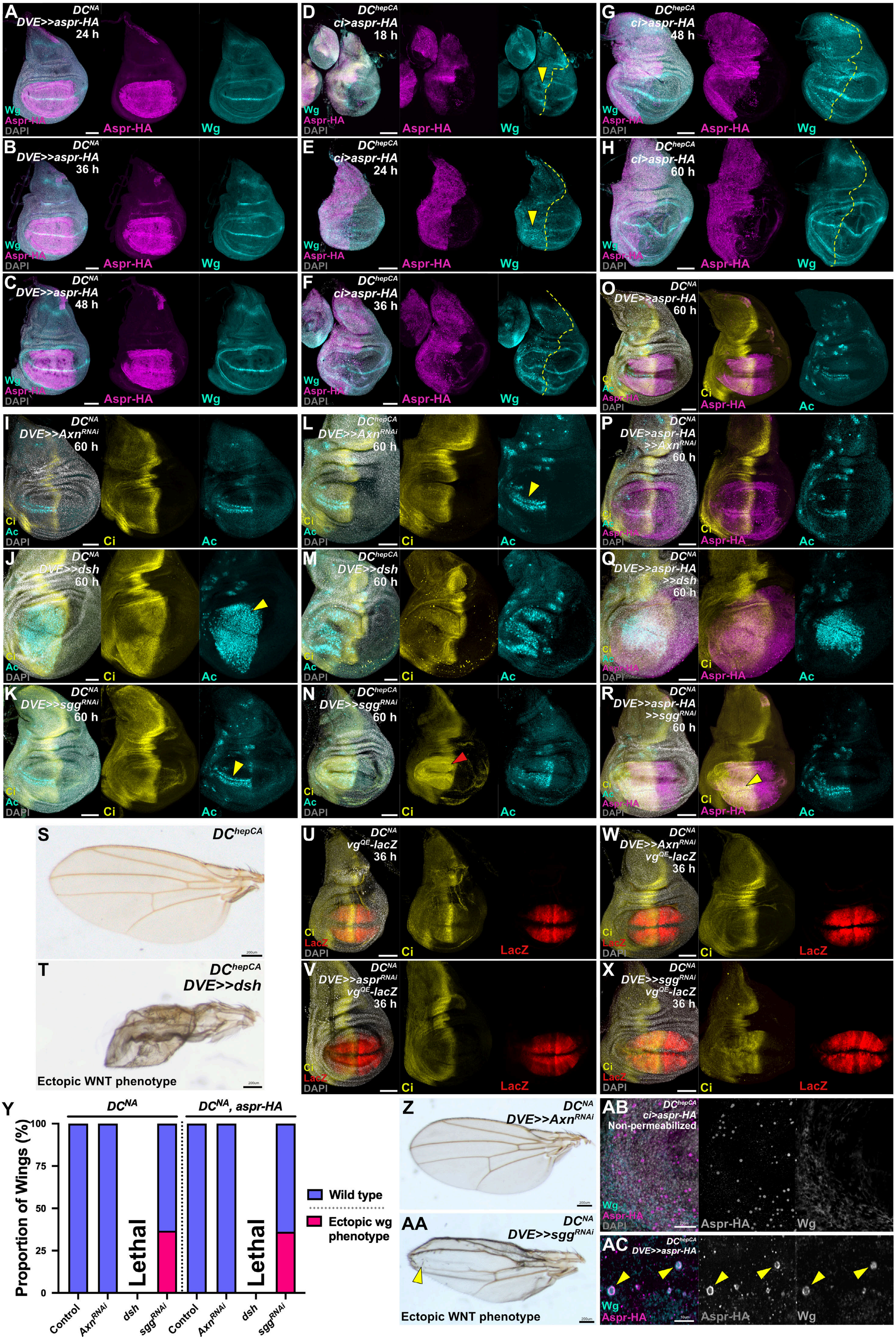

A DCNA DVE>>aspr-HA 24 h
B DCNA DVE>>aspr-HA 36 h
C DCNA DVE>>aspr-HA 48 h
D DChepCA ci>aspr-HA 18 h
E DChepCA ci>aspr-HA 24 h
F DChepCA ci>aspr-HA 36 h
G DChepCA ci>aspr-HA 48 h
H DChepCA ci>aspr-HA 60 h
Wg
Aspr-HA
DAPI
I DCNA DVE>>AxnRNAi 60 h
J DCNA DVE>>dsh 60 h
K DCNA DVE>>sggRNAi 60 h
L DChepCA DVE>>AxnRNAi 60 h
M DChepCA DVE>>dsh 60 h
N DChepCA DVE>>sggRNAi 60 h
O DCNA DVE>>aspr-HA 60 h
P DCNA DVE>>aspr-HA >>AxnRNAi 60 h
Q DCNA DVE>>aspr-HA >>dsh 60 h
R DCNA DVE>>aspr-HA >>sggRNAi 60 h
Ci
Ac
S DChepCA
T DChepCA DVE>>dsh
Ectopic WNT phenotype
U DCNA vgQE-lacZ 36 h
V DCNA DVE>>asprRNAi vgQE-lacZ 36 h
W DCNA DVE>>AxnRNAi vgQE-lacZ 36 h
X DCNA DVE>>sggRNAi vgQE-lacZ 36 h
LacZ
Y
DCNA
DCNA, aspr-HA
Proportion of Wings (%)
100
75
50
25
0
Lethal
Control
AxnRNAi
dsh
sggRNAi
Wild type
Ectopic wg phenotype
Z DCNA DVE>>AxnRNAi
AA DCNA DVE>>sggRNAi
Ectopic WNT phenotype
AB DChepCA ci>aspr-HA Non-permeabilized
AC DChepCA DVE>>aspr-HA

**Figure S9. Ectopic *Aspr* Modifies WNT Signaling – related to Figure 8**

(**A–C**) Unablated disc (*DC$^{NA}$*) at 24 h AHS expressing *aspr-HA* with *DVE>>GAL4* at 24–48 h post-ablation stained for Wg (cyan). Wg distribution remains unchanged in development.

(**D–H**) Ablated disc (*DC$^{hepCA}$*) expressing *aspr*-HA under *ci-GAL4* shows Wg condensing in the anterior compartment at 18–36 h (yellow arrowheads). Aspr-HA vesicles colocalize with Wg at all time points.

(**I–K**) Unablated disc (*DC$^{NA}$*) at 60 h AHS expressing WNT activation constructs (*Axn$^{RNAi}$, UAS-dsh, sgg$^{RNAi}$*) stained for Ci (yellow) and Ac (cyan)as a WNT target gene. *dsh* overexpression expands Ac expression (J), and *sgg$^{RNAi}$* thickens Ac (K).

(**L–N**) The same manipulations in (I-K) in ablated discs (*DC$^{hepCA}$*) similarly expand or thicken Ac expression and, with *sgg$^{RNAi}$*, increase Ci (red arrowhead).

(**O–R**) Co-expression of *aspr*-HA with WNT activation constructs as in (I–J) and (L–N) shows that *Aspr* weakens but does not abolish WNT-induced Ac phenotypes: incomplete Ac with *Axn$^{RNAi}$* (P), attenuated ectopic Ac with Dsh (Q), and moderated Ac expansion with *sgg$^{RNAi}$* plus increased Ci (yellow arrowhead) (R).

(**S**) Representative regenerated wing from ablated wild type discs (*DC$^{hepCA}$*).

(**T**) Representative regenerated wing from ablated discs (*DC$^{hepCA}$*) expressing Dsh shows severe malformations with characteristic ectopic WNT phenotype including patterning defects and ectopic bristles.

(**U–X**) *vgQE-lacZ* activity in unablated discs (*DC$^{NA}$*) expressing *aspr*$^{RNAi}$, Axn$^{RNAi}$, or sgg$^{RNAi}$ with *DVE>>GAL4* shows no change in reporter activation at 36 h AHS.

(**Y**) Quantification of ectopic WNT phenotype seen adult wings from undamaged discs (*DC$^{NA}$*) expressing WNT manipulations shown. Sample size: (*DC$^{NA}$*); control *y$^{RNAi}$* n = 158, *Axn$^{RNAi}$* n = 176, *sgg$^{RNAi}$* n = 207, (*DC$^{NA}$, aspr-HA*); control *y$^{RNAi}$* n = 120, *Axn$^{RNAi}$* n = 125, *sgg$^{RNAi}$* n = 94. Dsh overexpression in undamaged discs leads to pupal lethality.

(**Z**) Representative adult wing from unablated discs (*DC$^{NA}$*) expressing *Axn$^{RNAi}$* with *DVE>>GAL4*, showing no patterning defects or bristle changes.

(**AA**) Representative adult wing from unablated discs (*DC$^{NA}$*) expressing *sgg$^{RNAi}$* with *DVE>>GAL4,* showing characteristic ectopic WNT phenotype, including patterning defects and ectopic bristles.

(**AB**) High magnification view of ablated disc (*DC$^{hepCA}$*) expressing *aspr*-HA under *ci-GAL4* under non-permeabilizing conditions, stained for Aspr-HA (magenta) and Wg (cyan). DAPI (gray) shows cell nuclei. The Wg signal normally observed in Aspr-associated EVs is no longer detected in the absence permeabilization.

(**AC**) High magnification view of ablated disc (*DC$^{hepCA}$*) expressing *aspr*-HA under *DVE>>GAL4,* stained for Aspr-HA (magenta) and Wg (cyan). Wg is found both within and at the surface of the EV structures, where it overlaps Aspr-HA (yellow arrowheads).

Scale bars: 50 μm unless indicated. Full genotypes in Supplementary Genotypes.
